# pH-sensitive binder design with Proton-PottsMPNN

**DOI:** 10.64898/2026.09.30.755438

**Authors:** Christian P. Jacobsen, Lukas Deweid, Foster Birnbaum, Line Gormsen, Thomas Fryer, Ulla Jagd, Andreas Hald, Morten Rasch, Timothy Jenkins, Sergey Ovchinnikov

## Abstract

Physiological compartments span a wide pH range, from blood plasma near pH 7.4 through the mildly acidic tumor microenvironment to the acidified endosome and lysosome, and binders whose affinity responds to these differences underpin recycling antibodies, catalytic extracellular degraders and tumor-conditional therapeutics. Such binders are still obtained mainly by combinatorial histidine scanning and display selection, which is laborious and yields few transferable design rules. Recent computational approaches have enabled the design of pH-sensitive binders that engage at physiological pH and release under acidic conditions, by introducing histidines at key positions so that protonation generates electrostatic repulsion, evaluated through physics-based calculations. However, because the design model has no explicit representation of protonation state, it cannot shape the residues around a titratable site to favor one state over the other, leaving pH sensitivity to be screened for rather than specified. Here we present Proton-PottsMPNN, a sequence-design model that explicitly represents the protonated and deprotonated states of histidine, aspartate and glutamate as distinct sequence tokens. Because protonation state is specified rather than inferred, the model redesigns the microenvironment around a titratable residue so that the chosen protonation state is energetically favored over its alternative. Using Proton-PottsMPNN we designed a library of 8,407 *de novo* PD-L1 candidate binders in which sequence-structure compatibility and protonation state preference were designed jointly under a single energy function, with no structure-prediction filtering applied. The full library was synthesized and displayed on yeast, then screened against PD-L1 across a series of pH values by fluorescence- and magnetic-activated cell sorting, recovering 237 and 288 unique pH-dependent designs respectively. Transition pH values spanned 4.0 to 5.8, and switches were centered on protonated aspartate and glutamate as well as histidine. Recovered designs showed a stronger model preference for their designed protonation state than designs from the same backbone that were not recovered, and this preference was strongest in the designs that required the most acidic conditions to bind. Proton-PottsMPNN therefore designs binders that engage PD-L1 under acidic conditions fully de novo, directly from a specified protonation state.

## 1 Introduction

Protein binders encounter different pH environments depending on where they act or traffic: blood is near pH 7.4, the extracellular space of many solid tumours is around pH 6.0–6.5, and endosomal and lysosomal compartments range from pH 4.0 to 6.0 [1, 2]. These pH differences alter the protonation of titratable side chains, changing their charge and hydrogen-bonding capacity, which can in turn affect protein folding and the strength of interactions between binders and their targets. These effects have been exploited to improve the activity of protein drugs. Antibodies engineered to engage only at physiological pH and release their target in acidic environments have been shown to extend serum half-life through FcRn recycling [3, 4]. The same mechanism has been used to build catalytic LYTACs, which release their target in the late endosome and recycle to degrade further copies rather than being consumed one for one [5–7]. Engineering the opposite dependence, such that a binder engages only below pH 6.5, instead confines activity to the acidic tumour microenvironment and has been shown to widen the therapeutic window of conditionally activated antibodies and checkpoint inhibitors [8, 9]. Despite these applications, few methods generate pH-dependent binders by design [10, 11]. Most reported switches have been obtained by combinatorial histidine mutagenesis followed by phage or yeast display [12–14]. Although successful it is resource-intensive, gives little insight into why a given variant responds to pH, and findings transfer poorly between targets.

Ahn et al. recently showed that *de novo* design methods can also be used to design pH-sensitive binders. By biasing ProteinMPNN [15] towards histidine, placing histidines at the binder-target interface or in the binder core, and redesigning the surrounding residues, they generated binders that dissociate under acidic conditions and used them to build catalytic degraders [11]. pH responsiveness can therefore be engineered into designed binders, rather than screened for. The design model itself, however, has no explicit representation of protonation: ProteinMPNN can be biased towards histidine by composition, but it cannot distinguish the cationic form from the neutral one explicitly. Protonation enters only afterwards, through a Rosetta pose in which every histidine is doubly protonated [16] and scored by hydrogen-bond and electrostatic filters. This has three consequences. (i) Candidate sequences are filtered for pH sensitivity rather than explicitly designed for it, (ii) the local environment that sets a titratable residue’s pKa cannot be tuned directly, and (iii) the switch is restricted to histidine [11]. Histidine, however, is not the only titratable residue available. Buried or hydrogen-bonded aspartate and glutamate carboxylates can have pKa values raised by several units into the endosomal range [17, 18]. They also offer a mechanism histidine cannot: an anionic carboxylate placed near another negative charge is repelled in the deprotonated state, and protonation removes that repulsion, so the interaction is disfavored at neutral pH and permitted in acid [17]. Despite this, no current design method treats them as designable protonation states.

Here we introduce Proton-PottsMPNN which is an extension of the sequence-design model Potts-MPNN (Birnbaum and Keating [19]) with explicit protonation labels, which expands the standard 20-letter alphabet to 26 letters by assigning the titratable residues His, ASP and Glu with three tokens: two protonation state-specific tokens and one representing an undetermined protonation state (His/HIS-P/HIS-S, Asp/ASP-P/ASP-D, Glu/GLU-P/GLU-D; Methods section 4.2.). The protonation state is thus predicted together with residue identity, so the model can learn which local environments favor a protonated state. Like PottsMPNN, Proton-PottsMPNN was trained to produce two complementary outputs from a shared encoder: an autoregressive head, giving a per-position distribution over the expanded alphabet, and a Potts model head, giving single-residue and pairwise energy terms for the sequence as a whole; we evaluate both below, as they proved to capture different aspects of the sequence-structure relationship (Fig. **1**.a).

**Figure 1.**
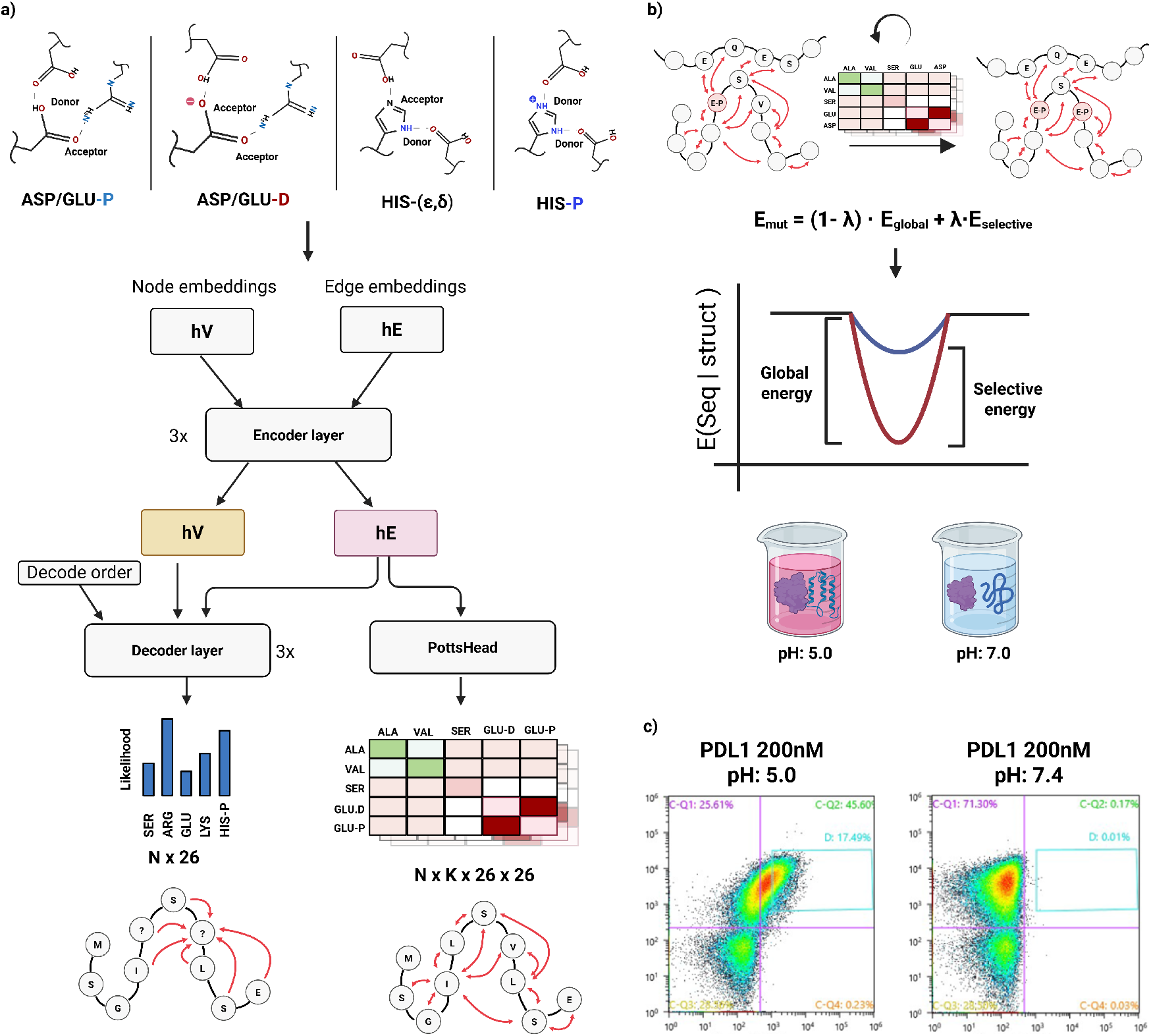
Proton-PottsMPNN and pH-conditioned binder design. **a)** Protonation-state prediction and model architecture. Local geometric features are used to predict protonation states, which are then encoded as sequence tokens for training Proton-PottsMPNN. ASP-P and GLU-P denote neutral, protonated carboxylic acids; ASP-D and GLU-D, deprotonated anionic carboxylates; HIS-S (*ε, δ*), single protonated neutral histidine; and HISP, cationic imidazolium. Backbone-derived node (*h_V_*) and edge (*h_E_*) embeddings are processed by shared encoder layers and passed to an autoregressive decoder and a Potts head, which produce token likelihoods and Potts energy terms, respectively. Red arrows represent graph interactions; *N* and *K* denote the numbers of residues and graph neighbors per residue. **b)** Protonation-conditioned redesign illustrated for a fixed ASP-P center (pink) relative to ASP-D (blue). *E*_D_ and *E*_P_ are the corresponding conditional Potts energies, and *E*_selective_ = *E*_P_ − *E*_D_. Neighboring residues are mutated to balance global Potts energy and protonation-state selectivity, weighted by *λ* where lower energies are more favorable. **c)** Flow cytometry of enriched binders from yeast-display library following incubation with 200nM biotinylated PD-L1 at pH 5.0 or 7.4.

We built on PottsMPNN because the Potts formulation yields more designable sequences and more accurate predictions of mutational ΔΔ*G* than likelihood-based scoring, which Birnbaum and Keating attribute to pair terms that represent constraints such as electrostatic repulsion explicitly [19]. Both properties are likely to be useful in pH-sensitive binder design. Because the encoder sees only backbone atoms, a single energy table scores every protonation assignment on a given backbone, so the preference between states and the stability of the fold are read out together. We find that the total Potts energy of a complex is predictive of designability, while the energy difference between protonation states correlates with measured p*K*_a_ for histidine, aspartate and glutamate. This lets us pose pH-sensitive design as a two-state problem: a set of titratable residues is pinned in a chosen protonation state and their local environments rewritten to favor that state over the alternative while remaining compatible with the backbone (Fig. **1**b).

Applying this framework, we designed a library of 8,407 *de novo* PD-L1 candidate binders around pinned protonated aspartate, glutamate and histidine, with sequence-structure compatibility and protonation state preference designed jointly under a single energy function and no post-hoc structure-prediction filtering applied. Fluorescence- and magnetic-activated cell sorting recovered hundreds of unique pH-dependent designs, with transitions spread across a pH series from 4.0 to 5.8 (Fig. **1**c). Recovered designs showed a stronger predicted preference for their designed protonation state than same-backbone designs that were not recovered, most strongly in those requiring the most acidic conditions. To our knowledge these are the first *de novo* designed binders for which acid-selective engagement was specified during design, on backbones not previously validated as binders: earlier computational work produced designed binders that release their target in acid [11], while binders that engage preferentially in acid have been obtained by histidine mutagenesis and display selection from antibodies already known to bind [8, 9, 14].

## 2 Results

### 2.1 A protonation-state-aware sequence-design model

Training a sequence-design model with explicit protonation state tokens requires a large set of protein structures in which the protonation labels are specified. However, experimental assignments are available for only relatively few structures. We therefore set out to train classifiers that predict protonation state from local structural environment, and to apply them across the Protein Data Bank (PDB). To obtain a training set we collected neutron and joint X-ray/neutron PDB structures with deposited H/D atom positions, reading the protonation state of each titratable residue directly from the experimentally determined proton positions. From the 216 collected structures we obtained 7,434 labeled residues: 416 HIS-P, 804 HIS-S, 36 ASP-P, 3,164 ASP-D, 58 GLU-P and 2,956 GLU-D. For each labeled residue we extracted 141 geometric features, including hydrogen-bond geometry, local density, neighborhood composition, and sequence context (Supp. Table 1). Using these features and the experimental labels, we trained XGBoost classifiers to predict protonation states using FLAML [20], and evaluated them by five-fold cross-validation stratified by structure, so that all residues from a given PDB entry fell in the same fold. Out-of-fold average precision was 0.70 for histidine and 0.33 for pooled acidic residues, corresponding to 2.1*×* and 21.9*×* over background (Supp. Fig. 1b). Before applying these classifiers across the PDB and AFDB to generate Proton-PottsMPNN training labels, we assessed whether the geometric environments present in the wider PDB were covered by those in the neutron-derived set. To this end, we compared the labeled residues with 10,262 histidines and 37,759 acidic residues sampled from the wider PDB and found broad overlap in feature space (Supp. Fig. 3).

Cross-validated performance is measured on the labeled residues themselves and says nothing about assignments made elsewhere. We therefore applied the classifiers to 1,000 PDB structures and asked whether the resulting labels were biophysically sensible. Across structural neighborhoods we find charge-consistent interactions, with HIS-P-assigned sites enriched for deprotonated acidic neighbors, HIS-S-assigned sites enriched for histidine neighbors, and protonated acidic sites enriched for protonated acidic neighbors (Supp. Fig. 1.f). Direct-coupling analysis of 1,070 multiple sequence alignments shows the same charge consistency, with HIS-P sites coupling significantly more strongly to acidic residues than HIS-S sites, and deprotonated Asp/Glu coupling significantly more strongly to basic residues than their protonated counterparts (Supp. Fig. 1.g). Predicted HIS-P, ASP-P and GLU-P states are also modestly but significantly enriched in lysosome-associated proteins relative to cytoplasm-associated ones (Supp. Fig. 1.e). Thus, labels assigned from local geometry alone recover charge-consistent relationships in both structural and evolutionary data.

We then applied the classifiers to histidine, aspartate and glutamate residues across 161,720 PDB and 199,843 AFDB structures (Methods, Sec. 4.2), obtaining a continuous protonation-state score for each residue. Converting these scores into protonation tokens requires a prediction cutoff, which defines the states the design model sees during training. To identify the best threshold we trained models over a range of histidine and pooled acidic cutoffs, scoring each against the experimental assignments in the H/D-resolved structures by negative log-likelihood of the reference token and by area under the precision-recall curve for protonation state prediction. Both criteria favored 0.3 for histidine and 0.06 for pooled acidic residues, giving average precision of 0.47 (1.6*×* over background) and 0.25 (16.6*×* over background) respectively (Fig. **2**.a). The upstream classifiers were trained on these same structures. This sweep therefore selects a cutoff rather than validating the labelling pipeline. The model trained at these cutoffs was used for all subsequent analyses. Its training set comprised *∼*125.4 million predicted titratable residues: 8.3 million HIS-P, 11.0 million HIS-S, 689,422 ASP-P, 48.4 million ASP-D, 942,628 GLU-P and 56.1 million GLU-D.

**Figure 2.**
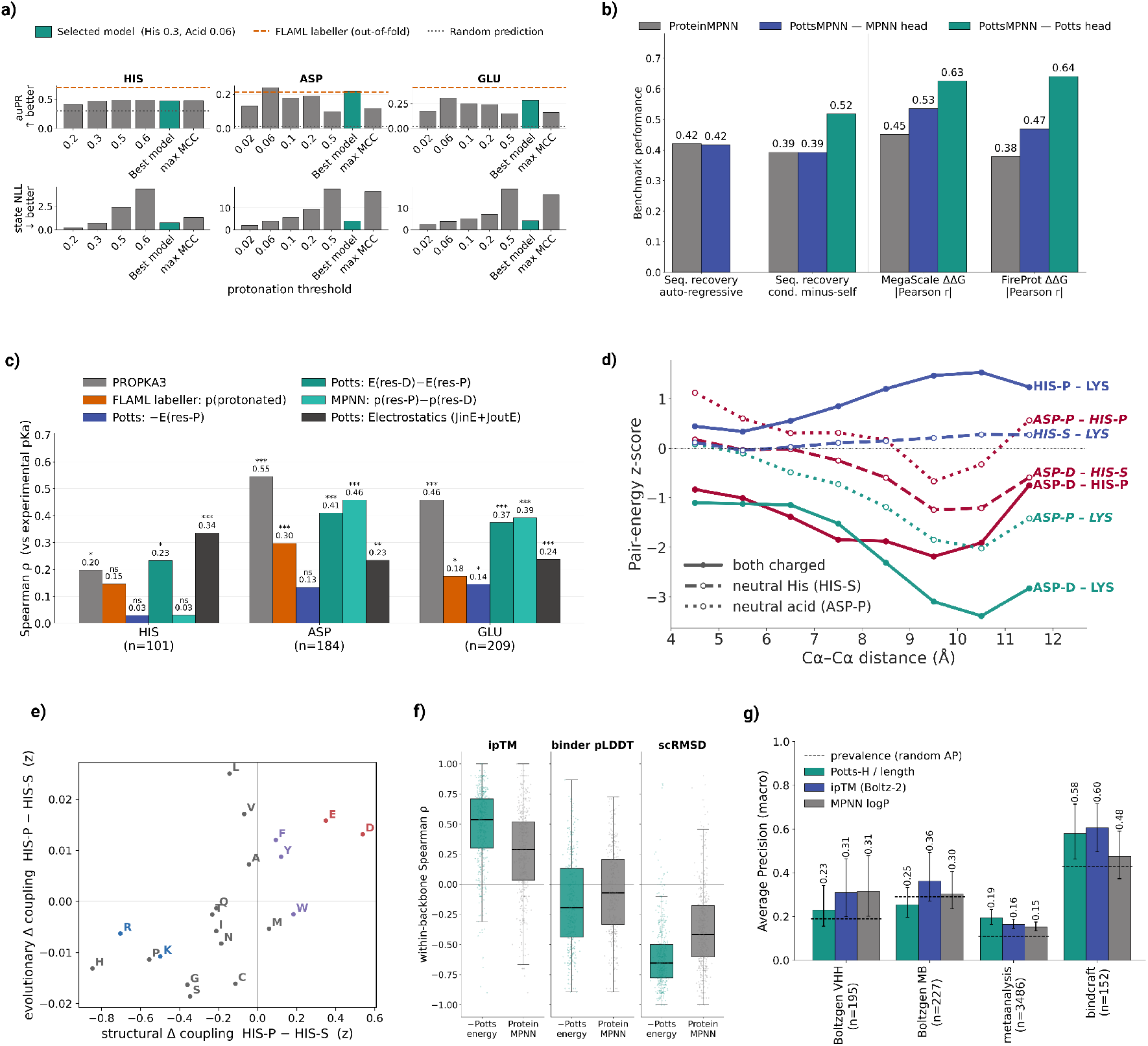
Model benchmarking and learned protonation-dependent interactions. **a)** Selection of protonation-label thresholds against experimentally assigned H/D states, for histidine (HIS), aspartate (ASP) and glutamate (GLU). Area under the precision-recall curve (top) and negative log-likelihood of the reference state (bottom) are shown per cutoff; teal marks the selected model (0.3 for HIS, 0.06 for the acids), the orange dashed line the upstream labelling classifier and the grey dotted line random prediction. **b)** Sequence recovery on 192 held-out chains under autoregressive and conditional-minus-self decoding, and absolute Pearson correlations with experimental ΔΔ*G*. Grey, blue and teal denote ProteinMPNN, the Proton-PottsMPNN decoder and its Potts head. **c)** Spearman correlations between experimental p*K*_a_ and PROPKA3 (grey) or model derived scores: the absolute protonated-state score (blue), the two-state gap from the Potts head (dark teal) and the decoder (light teal), and the gap restricted to electrostatic partners (black); *n* is the number of residues. Asterisks indicates two-sided *p*-values for *ρ ≠* 0 (* *p <* 0.05, ** *p <* 0.01, *** *p <* 0.001; ns, not significant). **d)** Mean standardized pair energies against C*α*– C*α* distance across 60 held-out structures, with line styles distinguishing charged pairs from those containing neutral HIS-S or ASP-P; negative values are more favorable. **e)** Standardised protonation-dependent change in structural couplings against the corresponding change in evolutionary couplings, for HIS-P relative to HIS-S. Points are the 20 partner amino acids, colored by charge. **f)**Within-backbone Spearman correlations between the negative Potts energy (teal) or ProteinMPNN decoder probability score (grey) and ipTM, binder pLDDT, or self-consistency RMSD (scRMSD) calculated based on RosettaFold3 [30]; each point is one backbone. **g)** Average precision for binder classification using length normalized Potts energy (teal), Boltz-2 ipTM (blue) or ProteinMPNN log-probability (grey) across four design classes. Dashed lines show binder prevalence, error bars are bootstrap confidence intervals from 2,000 resamples, and *n* is the number of designs.

### 2.2 Benchmarking Proton-PottsMPNN for sequence design and protonation awareness

Proton-PottsMPNN must do two things. It must design sequences that fold onto a given backbone, and it must distinguish protonation states in a way that reflects their real energetic preferences.

We first asked whether the model’s preference for protonated over deprotonated states tracks experimental p*K*_a_. Since p*K*_a_ reflects the environment-dependent balance between the two states, we computed their likelihood difference from the autoregressive head and their energy gap from the Potts head (Methods). Neither score is calibrated in p*K*_a_ units, so we used Spearman rank correlation against the PKAD database [21], covering 101 histidines, 184 aspartates and 209 glutamates from 106 structures, with PROPKA3 as a benchmark [22] (Fig. **2**.c). We found that the two-state Potts gap correlated significantly with experimental p*K*_a_ for all three residue types (His *ρ* = 0.23, *p* = 0.019; Asp *ρ* = 0.41, *p* = 7 *×* 10*^−^*^9^; Glu *ρ* = 0.37, *p* = 2 *×* 10*^−^*^8^; Fig. **2**.c). For histidine it was comparable to PROPKA3 (*ρ* = 0.20, *p* = 0.047) and improved even further when the score was restricted to pairwise terms involving charged partners (*ρ* = 0.34, *p* = 6 *×* 10*^−^*^4^). Still, PROPKA3 remained the stronger predictor for the acidic residues (Asp *ρ* = 0.55; Glu *ρ* = 0.46). Although the autoregressive scores correlated significantly with p*K*_a_ for aspartate (*ρ* = 0.46, *p* = 6 *×* 10*^−^*^11^) and glutamate (*ρ* = 0.39, *p* = 4 *×* 10*^−^*^9^) it showed no predictive power for histidine (*ρ* = 0.03, *p* = 0.77). Furthermore, we found that the absolute protonated-state score was at best weakly predictive of p*K*_a_ (His *ρ* = 0.03, *p* = 0.77; Asp *ρ* = 0.13, *p* = 0.070; Glu *ρ* = 0.14, *p* = 0.037), and the signal thus appeared in the difference between the two states (*ρ* = 0.03 versus *ρ* = 0.23 for histidine; Fig. **2**.c). The model therefore ranks p*K*_a_ by the relative preference between protonation states rather than by the likelihood of the protonated state alone. Interestingly, the XGBoost probabilities from which the training labels were derived did not significantly rank histidine p*K*_a_ on the same residues (*ρ* = 0.15, *p* = 0.14), and ranked the acidic residues less well than the Potts gap (Asp *ρ* = 0.30; Glu *ρ* = 0.18; Fig. **2**.c).

We then examined if the pairwise terms from the learned Potts model discriminated between protonation states. To this end, we scored alternative protonation-state pairs on side-chain-facing edges from 60 PDB structures held out during training. Because the encoder depends only on the backbone, every token pair can be evaluated at every edge independently of the residue’s actual identity. We found that normalized pair energies showed the expected Coulomb-like distance dependence, with like-charge pairs penalized at short range and opposite-charge pairs favored, as also seen by Birnbaum and Keating [19] (Fig. **2** d). We observed that these preferences shifted with protonation state: protonating histidine made interactions with Lys less favorable and those with deprotonated Asp and Glu more favorable, whereas deprotonating Asp or Glu reversed the pattern. Comparatively the neutral HIS-S and ASP-P tokens fell between the charged extremes. The same contrasts were observed in evolutionary data: per-partner coupling changes upon protonation correlated with those inferred by direct-coupling analysis over 1,070 MSA alignments (Pearson *r* = 0.62, *p* = 4 *×* 10*^−^*^3^ for histidine, *r* = 0.6, *p* = 5 *×* 10*^−^*^3^ for aspartate, *r* = 0.67, *p* = 1 *×* 10*^−^*^3^ for glutamate; Fig. **2**.e), with the largest shifts at charged and polar partners. Thus, the Potts head seems to represent the formal charge of each protonation state and carries it into its pairwise energies.

We also found that expanding the alphabet did not compromise conventional sequence-design performance. On the held-out PDB and AFDB validation set the autoregressive head reached a canonical sequence recovery of 0.44, and on a 192-structure subset both Proton-PottsMPNN and ProteinMPNN recovered 0.42 under autoregressive decoding. Furthermore, we found that the Potts energy correlated more strongly with measured stability changes than the likelihood-based scores from either decoder (Fig. **2**.b) on the MegaScale [23] and FireProt [24] mutational-stability sets. These results reproduce the advantage Birnbaum and Keating reported for pairwise supervision, now over an expanded vocabulary.

Because the intended application of Proton-PottsMPNN is the design of pH-responsive binders, we next asked whether the Potts energy carried information about protein-protein binding. We first asked whether the energy could rank designs within a fixed backbone, using 10,469 ProteinMPNN-designed sequences across 463 RFdiffusion-3 [25] backbones spanning 11 target chains from the validation set. For each backbone we correlated the Potts energy with ipTM, binder pLDDT and self-consistency RMSD (scRMSD), taking the ProteinMPNN decoder probability score on the same designs as a baseline. Across most of the backbones, lower Potts energies accompanied higher ipTM and lower scRMSD (median *ρ* = 0.55 and *ρ* = *−*0.67), with correlations stronger than the ProteinMPNN baseline (Fig. **2**.f). Ranking by Potts energy, or minimising it during design, could therefore serve as an inexpensive pre-filter ahead of folding-based structure prediction.

We then asked whether the model separated experimentally confirmed binders from non-binders, using 4,060 designs from a meta-analysis dataset [26], BindCraft designs [27], and BoltzGen minibinders and nanobodies [28]. Here we scored complexes by total Potts energy divided by the number of residues, which removes the length dependence of the unnormalized energy across designs of differing size. Complexes were represented by Boltz-2 [29] predicted structures, and ipTM from the same predictions served as the baseline, so the two scores differ in scoring rather than input. The Potts energy showed competitive predictive performance on the meta-analysis dataset (AP 0.19 versus 0.16), and BindCraft dataset (0.58 versus 0.60), and fell below it on both BoltzGen classes (0.25 versus 0.36 for mini-binders; 0.23 versus 0.31 for nanobodies; Fig. **2**.g). Sequence-structure compatibility is therefore predictive of binding, though it does not consistently outperform a structure-based confidence metric.

Together, these results show that Proton-PottsMPNN retained the general sequence-design and stability-prediction capabilities of PottsMPNN trained by Birnbaum and Keating while learning an explicit representation of residue protonation. Its protonation-state scores were associated with experimental p*K_a_* values, and its pairwise energy terms agreed with protonation-dependent coupling patterns observed in evolutionary data and aligned with known electrostatic effects. The model also produced a relative binding score that discriminates binding/non-binding designs and can be used as cheap pre-filter for selecting designs, supporting its use for the design and evaluation of pH-responsive protein interactions.

### 2.3 Using Proton-PottsMPNN for multi-state design

Having established that Proton-PottsMPNN recovers experimentally assigned protonation states and ranks residues by measured p*K*_a_, we asked whether it could design sequence environments that favor specified protonation states. We formulated this as a two-stage workflow. First, one or more titratable residues are pinned to a target protonation-state token, either at user-defined positions or at sites ranked by a model-guided placement scan (Methods section 4.4). Second, residues surrounding each pinned titratable center are redesigned against two objectives: the selective energy, the gap between the target protonation state and its alternative, and the global Potts energy of the design, with a weight *λ* setting the balance between the objectives (Methods section 4.4).

We benchmarked the two redesign strategies across *λ ∈ {*0, 0.1*,…,* 1.0*}* on 37 seed binders. These seed binders were generated by RFdiffusion-3 [25] backbone design followed by ProteinMPNN sequence design against eight targets drawn at random from the PDB validation set, and retained only if RosettaFold3 predictions gave ipTM *>* 0.8, pLDDT *>* 80 and scRMSD *<* 2.5 Å. We evaluated performance of pH sensitive design on three challenges: neutral histidine for a high-pH target state, and cationic histidine and the neutral acidic states for a low-pH target state.

For all three tasks, MCMC sampling on the Potts energies gave greater control over the selective-energy gap than the autoregressive decoder, which reached a limited range of selectivity regardless of weight (Fig. **3.a**, Supp. Fig. 6.c). This motivated a hybrid strategy in which the autoregressive head generated infills around the fixed titratable centers and the Potts head scored the resulting sequences for their preference between the target and alternative states. Relative to decoder-only infilling, this gave controllable decreases in selective energy (Supp. Fig. 6.d; Methods, Sec. 4.4). We found that the two design objectives traced a Pareto front, with the lowest selective energies carrying the largest penalty in global Potts energy relative to the seed and producing less designable sequences, as reflected in scRMSD and ΔipTM (Fig. **3**.a). Selecting designs along the front nonetheless recovered sequences combining low selective energy with high predicted recovery of the target protonation state (microstate match), limited loss of interaction confidence, and self-consistency comparable to the seeds (Fig. **3**.c). The mechanism by which the selective gap was widened differed by target protonation state. Increasing *λ* built stabilising salt bridges for cationic histidine, and destabilising charge repulsion for neutral histidine and the acidic states (Fig. **3**.a, Supp. Fig. 7).

**Figure 3.**
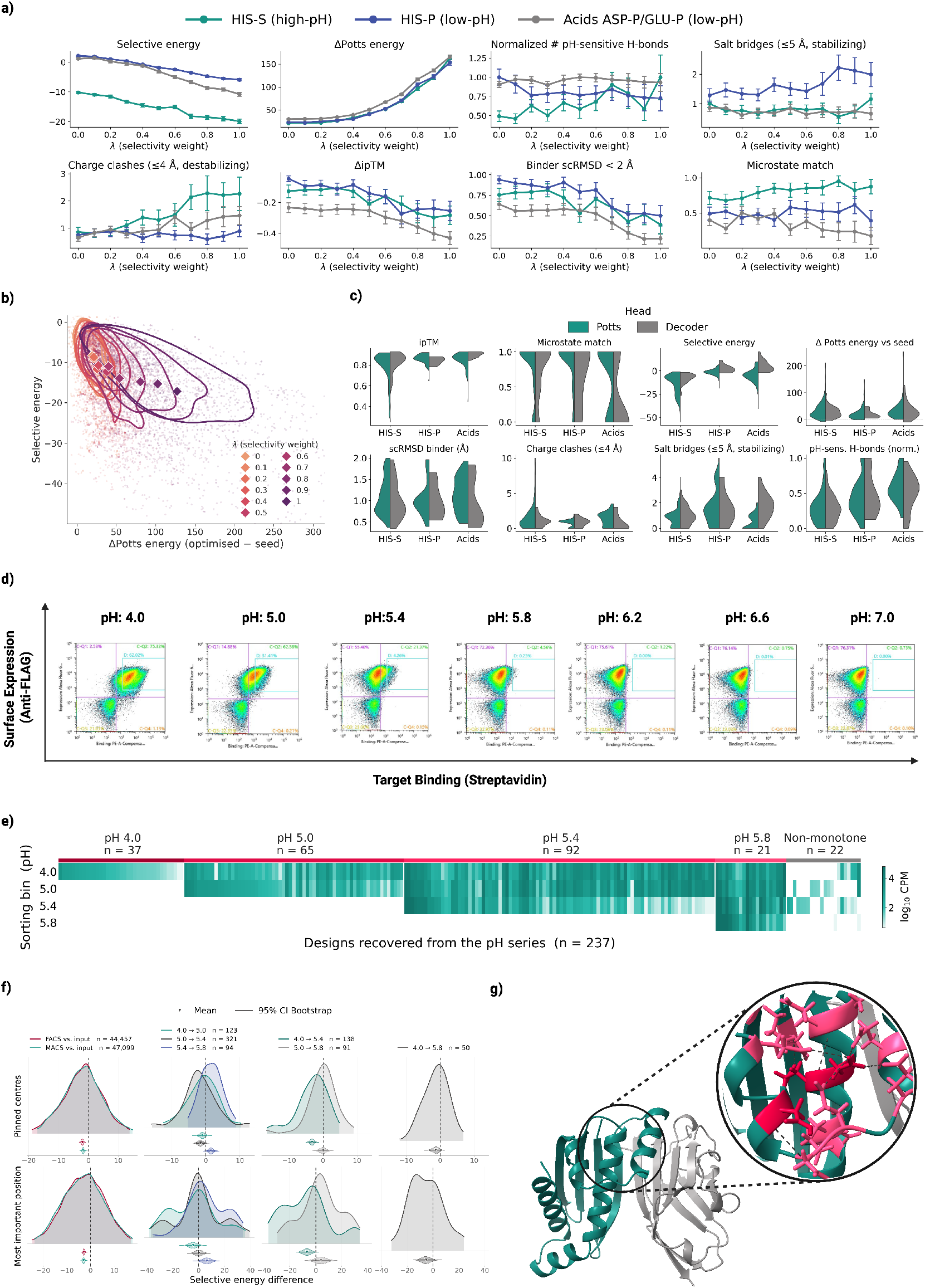
Benchmarking pH-conditioned design and characterizing PD-L1 binders. **a)** Effect of the selectivity weight *λ* on energetic, sequence and structural metrics for 3,183 RosettaFold3-refolded designs optimized with the Potts energy, from 37 seed binders across eight targets (decoder counterpart, Supp. Fig. 6c). Teal, blue and grey denote HIS-S (high-pH target state), HIS-P and pooled ASP-P/GLU-P (low-pH) designs. *λ* = 0 weights the global Potts energy alone, *λ* = 1 the selective energy alone; lower selective energy indicates a stronger preference for the target state. Δ values are relative to the seed; error bars, standard error across designs. **b)** Selective energy against the change in Potts energy from seed to optimized sequence, colored by *λ*; diamonds give the median per *λ* and contours the design density around it. **c)** Split violins of structural, energetic and sequence properties for Pareto-selected designs generated with the Potts energy (teal) or the autoregressive decoder (grey), shown per design challenge. **d)** Flow cytometry of the enriched library at the indicated pH following incubation with 200 nM biotinylated PD-L1; percentages give the fraction of events in each region. **e)**Sequencing abundance (log10 CPM) of 237 recovered designs across the pH-sorting pools, grouped by the highest pH at which binding was retained; non-monotonic profiles shown separately. **f)** Selective-energy differences between matched same-backbone designs, across all pinned centers (top) or at the most important position by Δ*E* (bottom), taken as recovered (or higher-pH) minus other. Leftmost column, recovered versus unrecovered designs for the FACS and MACS tracks; remaining columns, designs assigned to the indicated pH-response bins. Curves, paired differences; shaded distributions, bootstrap estimates of the mean; bars, 95% confidence intervals; *n*, matched pairs. **g)** Predicted structure of an example recovered binder (teal) with PD-L1 (grey); pinned residues in red, redesigned surrounding positions in pink.

We next applied the framework to generate PD-L1 binders intended to bind preferentially under acidic conditions. As before, seed binders were generated by RFdiffusion-3 and ProteinMPNN and filtered with RosettaFold3 against four PD-L1 epitopes, yielding 192 seeds. From these we generated 172,000 candidate variants carrying one to five pinned titratable sites assigned to HIS-P, ASP-P or GLU-P, placed at the interface or in the binder core either at random or by the model-guided placement scan. Guided by the benchmark, we applied both redesign strategies at *λ ∈ {*0.3, 0.4, 0.5*}* and selected 8,215 sequences by sampling across selective and global Potts energy scores, omitting post-redesign structure-prediction filtering because current models may not reliably resolve the geometries of protonation-dependent interactions.

These 8,215 designs and 192 ProteinMPNN controls were prepared as a yeast-display library and screened by two routes. In the first, three rounds of fluorescence-activated sorting (FACS) at pH 5.0 enriched the library for PD-L1 binding, collecting both an inclusive and a stringent gate at each round. The inclusive population was then sorted a fourth time into pH bins between 4.0 and 7.4. Because FACS sheath fluid is held at physiological pH, this route could miss binders whose low-pH complexes dissociate during the seconds spent at neutral pH before detection. We therefore ran a second selection route in which four rounds of magnetic-activated cell sorting (MACS) maintained acidic conditions from capture through washing, with elution at pH 7.4. All sorted pools, the initial library and a population sorted for expression alone were sequenced (Methods, Sec. 4.5).

We found that both the inclusive and the stringent sorting gates yielded populations that bound PD-L1 at pH 5.0 but not at pH 7.4 (Supp. Fig. 10). Counter-screening the enriched population at pH 5.0 against two unrelated proteins, HSA and CD98, showed that this binding was largely target-specific (Supp. Fig. 12).

FACS sorting the inclusive track into pH bins resolved binding populations at pH 4.0 and 5.0, smaller populations at pH 5.4 and 5.8, and none at pH 6.2 (Fig. **3**.d). Sequencing of these pH bins recovered 237 unique designs, *∼*3% of the library. Assigning each to the highest pH at which it remained detectable, 37 (16%) were lost above pH 4.0, 65 (27%) above pH 5.0 and 92 (39%) above pH 5.4, while 21 (9%) were retained at pH 5.8 (Fig. **3**.e). The remaining 22 designs (9%) appeared in non-consecutive bins, could not be assigned a single transition pH, and are shown separately. Most binders therefore showed monotonic profiles, remaining detectable across consecutive bins before dropping out at a characteristic pH, though the loss was sharp for some designs and gradual for others (Supp. Fig. 14.b). The recovered designs were concentrated on a small number of backbones, within which sequences showed different pH-sensitive profiles (Supp. Fig. 14.a). A small fraction of the ProteinMPNN seeds (0.5%) was recovered in the enriched population after three rounds of sorting, but none appeared in any pH bin (Supp. Fig. 11).

MACS recovered 288 binders after four rounds, of which a large population overlapped with the FACS set, indicating that the neutral interval before interrogation does not systematically discard pH-sensitive binders (Supp. Fig. 15). The MACS recoveries were enriched for protonation-conditioned designs but still contained a small subpopulation of the original ProteinMPNN seeds. Because few of the original ProteinMPNN designs were recovered at physiological pH, we ran a second sorting experiment at pH 7.4 with the target tetramerised and at raised concentration, so that weaker binders could be captured through avidity. This recovered 1.5% of the ProteinMPNN designs against 1.3% of the protonation-conditioned designs, which showed no binding under the pH 5.0 conditions (Supp. Fig. 11.d).

We then compared the scores of the recovered designs with the input library to identify which design properties were associated with pH-dependent binding. The two design objectives, the selective energy and the global Potts energy, are both sums over a backbone’s own fields and couplings, so their absolute values depend on chain length, contact density and local geometry, and comparisons across unrelated backbones are dominated by those differences rather than by the pinned centers. Pooled across the library, the selective energy nonetheless shifted in the expected direction over successive rounds of magnetic selection and across the FACS pH series, while the global Potts energy was essentially unchanged in both tracks (Supp. Fig. 13.a-c; Supp. Fig. 16.a-c). These shifts are small relative to the spread within each pool, so neither score separates recovered from unrecovered designs significantly, though the direction of the selective-energy shift is consistent across both selection routes. We therefore restricted all subsequent energy comparisons to designs sharing a backbone.

Across 44,457 matched pairs spanning 53 backbones, recovered designs had significantly lower protonation-selective energies than designs from the same backbone that were not recovered (mean difference *−*2.03, 95% CI [*−*2.69*, −*1.37] for FACS and *−*1.78, 95% CI [*−*2.39*, −*1.01] for MACS; Fig. **3**.f). The same relationship held within the pH series. Designs that lost binding at lower pH had generally lower selective energies than same-backbone designs that retained binding at higher pH, both between the pH 4.0 and 5.4 bins and between the pH 5.0 and 5.4 bins, with significance assessed by bootstrap resampling of matched design pairs (Fig. **3**.f). The same trend appeared between pH 4.0 and 5.0 and between pH 4.0 and 5.8 without reaching significance.

Design composition, unlike the energies, is comparable across the whole library. The HIS-P share fell from 0.39 in the input library to between 0.25 and 0.28 across the pH bins, with a corresponding rise in ASP-P, and designs carrying a single pinned center were strongly depleted (0.17 to 0.04), whereas the distribution of *λ* was essentially unchanged (Supp. Fig. 13.f-g).

Recovered designs carried more pH-sensitive hydrogen bonds at the pinned centers and more destabilising charge repulsion than unrecovered same-backbone designs, while the number of salt bridges was lower (Supp. Fig. 17). By contrast, PROPKA3 p*K*_a_ values at the pinned centers separated neither recovered from unrecovered designs nor designs in adjacent pH bins (Supp. Fig. 18), and no structure-prediction metric computed after sequencing was predictive of binding at lower pH (Supp. Fig. 13.h-j; Supp. Fig. 18.h-j).

## 3 Discussion

Proton-PottsMPNN makes the protonation states of histidine, aspartate and glutamate explicit during sequence design. Its selective energy gap between protonation states correlates significantly with experimental p*K*_a_ for all three residue types, whereas the protonated-state score alone gives a weaker and less consistent signal (Fig. **2**.c), indicating that the model ranks p*K*_a_ by the relative preference between states rather than by the likelihood of the protonated one. For histidine the correlation is numerically higher than that of PROPKA3 and rises when the gap is restricted to charged partners, while PROPKA3 is stronger for the acidic residues, though paired comparisons established no significant difference for any residue type. The gap is nonetheless a model preference conditional on the assigned neighboring states, not a p*K*_a_ in physical units or a prediction of binding transition pH. The learned pairwise energies distinguish the states through their neighbors: protonated histidine is less compatible with nearby cations and more compatible with anions than neutral histidine, and the acidic residues show the opposite preference between their states (Fig. **2**.d; Supp. Fig. 7). These preferences vary with distance and broadly agree with evolutionary couplings (Fig. **2**.e; Supp. Fig. 5.c), consistent with charge-sensitive interactions.

The practical advantage of the Potts formulation for design is that one energy function scores both the selective energy gap at a pinned site and the sequence-wide cost of the substitutions made around it. An autoregressive decoder can compare state-dependent preferences locally but cannot measure that cost, and in the design benchmark direct Potts optimization gave considerably greater control over the selective energy gap (Fig. **3**.a-c; Supp. Fig. 6.c-d). We find that the global Potts energy and the selective energy gap trace a Pareto front, with more selective designs showing worse global energy scores and structural self-consistency. Still, selecting along that front gave designs that were both designable and protonation-selective (Fig. **3**.b-c).

Using a two-state design approach, we redesigned 192 RFdiffusion-3 and ProteinMPNN seed binders into 8,407 *de novo* PD-L1 candidates. Mutations were introduced either by Potts optimization, which balanced the global Potts energy against the selective energy gap, or by hybrid autoregressive infilling, which combined decoder probabilities with the same selective energy gap from the Potts head. Although the seed binders had passed structural filters (ipTM *>* 0.8, binder pLDDT *>* 80, binder scRMSD *<* 2.5 Å), they had never been experimentally validated as binders, and no structure-prediction filtering was applied after redesign. These candidates were then screened by two sorting routes, FACS and MACS, each of which tests binding and pH sensitivity in the same experiment. The recovered pH-dependent binders were concentrated on a small number of backbones, and counter-screening showed binding at pH 5.0 to be largely target-directed (Fig. **3**.d-e; Supp. Fig. 12). The unmodified seeds, included as controls, were recovered after the second FACS round but in none of the pH bins, and although MACS also enriched them, it did so less than the conditioned designs. A separate selection at pH 7.4 with the target concentrated and tetramerised recovered the seeds at a slightly higher rate than the pH-conditioned designs (1.5% against 1.3%), consistent with the conditioned designs having traded neutral-pH engagement for acid selectivity, though the margin is small.

To our knowledge these are the first *de novo* designed binders for which acid-selective engagement was specified during design. Ahn et al. designed binders that release their target in acid and proposed the converse as an extension of their method, while binders that engage preferentially in acid have otherwise come from histidine mutagenesis and display selection of existing antibodies [8, 9, 11, 14]. Supporting the intended mechanism, the selective energy gap at the pinned centers predicted recovery within a backbone while the global Potts energy did not (Fig. **3**.f), indicating that the designed preference for a protonation state, rather than general designability, tracks acid-dependent binding. PROPKA3 p*K*_a_ at those centers was likewise unpredictive (Supp. Fig. 18), so the signal is not reproduced by a static electrostatic estimate on the predicted complexes. Still, the designs are so far pH-dependent only at the level of display enrichment. Affinities at each pH on purified protein remain to be measured, and establishing that the pinned residues are themselves responsible will require mutating them back in individual designs.

Protonation-state assignments across the PDB and AFDB training collections were classifier-derived, and the classifiers themselves were trained on only 216 H/D-resolved structures containing 416 cationic histidines and 94 protonated aspartates and glutamates, the latter just 1.5% of the labeled acidic residues. The model therefore learned from predicted labels at a scale four orders of magnitude beyond the experimental evidence behind them. At the selected cutoffs it recovers the assigned states only moderately, reaching an average precision of 0.47 for histidine and 0.25 for the acids, in both cases below the labelling classifiers themselves (Fig. **2**.a). Part of this difference is expected, since the classifiers read side-chain geometry while the model conditions on the backbone alone, but it means the state tokens are better read as relative preferences than as state assignments.

Despite this, Proton-PottsMPNN ranked experimental p*K*_a_ significantly for aspartate and glutamate as well as histidine, learned pairwise energies consistent with the formal charge of each state, and produced acid-centered switches that survived selection, with ASP-P in fact enriched among recovered designs relative to the input library. Weak supervision at the level of individual labels was therefore sufficient to place the states correctly relative to one another, which is what the two-state objective depends on. Agreement with MSA-derived couplings supports this, though it is a consistency check rather than a fully independent one, since coevolutionary couplings can reflect residue contacts and local contact geometry and charge composition was used by the classifier-derived labels. Better labels should nonetheless help, and how they are obtained matters. Assigning states from hydrogen-bond and salt-bridge geometry alone gave weaker p*K*_a_ ranking and weaker discrimination between recovered and unrecovered designs (data not shown), so simple geometric rules are not a substitute for learned assignments. Additional H/D-resolved structures or carefully validated constant-pH simulations are therefore an alternative, and improved labels are the change most likely to improve the model.

Several extensions follow from the framework. Birnbaum and Keating obtained their strongest models using multiple sequence alignments, so incorporating evolutionary information while retaining explicit protonation tokens may improve state assignment and protein design further [19]. In this work we modeled protonation explicitly only for histidine, aspartate and glutamate, because the study aimed at designing binders that engage under acidic conditions, but distinct charged and neutral lysine tokens would extend the approach to other pH-dependent problems. Beyond binders, redesigning the environments of catalytic titratable residues could test whether a change in protonation-state preference alters an enzyme’s pH-activity profile. Furthermore, ProtonPottsMPNN conditions on a given structure, so it can rewrite the sequence around a titratable site but cannot change the backbone in response to protonation. Adding protonation tokens to models that generate or reason over structures would make that response designable, which is likely what pH-dependent conformational and fold switches require.

Conditioning sequence design on protonation states thus yielded binders that engage PD-L1 under acidic conditions from fully *de novo* backbones in FACS and MACS. We found that the selective energy gap between protonation states both ranked experimental p*K*_a_ and separated recovered from unrecovered designs within a backbone. We are currently testing the inverse response, redesigning existing binders to release their target under acidic conditions, which asks whether the same framework controls the switch in both directions.

## 4 Materials and Methods

### 4.1 Protonation-state labeling

Protonation-state prediction was formulated as two binary-classification tasks. The histidine classifier distinguished the cationic imidazolium state from the singly protonated, neutral imidazole state. The acid classifier was trained jointly on aspartate and glutamate and distinguished the neutral carboxylic-acid state from the deprotonated, anionic carboxylate state.

Reference labels were taken from neutron and joint X-ray/neutron PDB structures with deposited H/D atom positions, assigned as described in Section 4.3. For each labeled residue we computed 138 (histidine) or 141 (acid) geometric features spanning burial and density, neighborhood composition, distances to chemical groups, hydrogen-bond roles and geometry, an a-priori electrostatic field, and sequence context (Supp. Table 1). Hydrogen bonds were identified with HBPLUS [31] (version 3.2) under permissive criteria (*d*_HA_ *≤* 3.2 Å, *d*_DA_ *≤* 4.0 Å, *θ*_DHA_ *≥* 90*^◦^*). Because the default HBPLUS donor list excludes carboxyl oxygens, bonds were enumerated in three tagged passes and merged for aspartate and glutamate (Supp. Methods). All deposited hydrogen atoms were removed before annotation, and features were computed from protein heavy atoms only.

Classifiers were trained using FLAML[20] under grouped five-fold cross-validation, with observations grouped by PDB identifier so that all residues from a structure fell in the same fold. Models were selected by mean average precision (AP) across folds, treating the protonated state as the positive class, because of the large class imbalance between protonated and deprotonated states, particularly for the acidic residues. Out-of-fold probabilities were concatenated across folds to construct precision-recall curves and the reported AP values. Each classifier was then refitted on all labeled observations and applied to histidine, aspartate and glutamate residues in the PDB and AFDB structure sets. A rule-based alternative, assigning states directly from HBPLUS donor and acceptor patterns under stricter geometric criteria, is described in Supp. Methods and shown for comparison in Supp. Fig. 1.

### 4.2 Model and training

#### Extended sequence vocabulary

The vocabulary comprised the 20 canonical amino-acid tokens, one unknown token, and nine protonation-state tokens, three per titratable residue type, giving *V* = 26 tokens (*V* = 30 including ambigous tokens). The suffix P denotes the protonated state, cationic imidazolium for histidine and neutral carboxylic acid for the acids; S denotes the singly protonated, neutral imidazole; D a deprotonated, anionic carboxylate; and A an ambiguous classifier assignment rather than a distinct biochemical state.

Each titratable residue was assigned one token using the five-fold classifier ensemble, taking the A token when the ensemble standard deviation exceeded 0.20 (histidine) or 0.30 (acids). The canonical parent tokens were retained for label dropout: on each training presentation every titratable residue was represented independently by its protonation-state token with probability 0.5 and by its parent token otherwise, applied identically to the target and the teacher-forced context. Validation used the assigned tokens without dropout. Neutral histidine was represented by a single HIS-S token rather than separate tautomers, because the model receives only backbone heavy atoms and cannot infer which ring nitrogen is protonated. A tokens were supervised as prediction targets; only UNK tokens were excluded from the loss.

#### Model

Proton-PottsMPNN follows the PottsMPNN framework [19], a ProteinMPNN-style graph encoder-decoder operating on backbone heavy atoms. An autoregressive decoder predicts a distribution 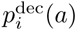 over the *V* tokens at each position *i* in a random decoding order. A second head predicts single-site fields *h_i_*(*a*) and pairwise couplings *J_ij_*(*a, b*), which for a token sequence *s* of length *L* define the Potts Hamiltonian

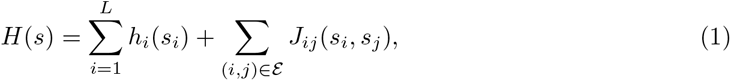

where *ε* is the directed edge set and lower values are more favorable. Because the fields and couplings depend only on the backbone, the Hamiltonian scores alternative token sequences, including alternative protonation assignments, without further encoder passes. Relative to the reference implementation the vocabulary was increased to *V* = 26 and reciprocal-edge couplings were symmetrized by averaging the two directions.

The conditional energy of a candidate token *a* at position *i*, with all other positions fixed, was

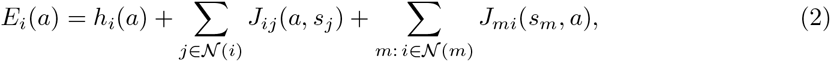

giving *P_i_*(*a*) = exp[*−E_i_*(*a*)]*/* Σ*_a_*_′_ exp[*−E_i_*(*a′*)]. Both coupling sums are required because the neighbor graph is directed and approximately 16% of edges lack a reciprocal partner.

#### Training data and objective

Training examples were individual polymer chains from the first biological assembly of PDB entries, determined by X-ray diffraction or electron microscopy at better than 3.5 Å, containing 30-10,000 non-atomized tokens and deposited before 2022-12-16; NMR structures were excluded. Sequences were clustered with MMseqs2[32] at 40% identity and 60% coverage, and 10% of clusters were held out for validation. The set was supplemented with AFDB[33] dimers passing ipTM *≥* 0.80, pDockQ2[34] *>* 0.23 and mean interface pLDDT *>* 85, clustered in a common namespace with the experimental chains to keep the splits cluster-disjoint (Supp. Methods). Each batch contained 50% experimental chains, 25% predicted homodimers and 25% predicted heterodimers, sampled within each source with probability proportional to *n*_prot_*/N*_clust_. Examples were cropped to the query chain and its contacting chains subject to a 6,000 heavy-atom limit, and isotropic Gaussian coordinate noise of standard deviation 0.2 Å was resampled on every presentation during training only.

The two heads were trained jointly with equally weighted losses: label-smoothed cross-entropy (*ε* = 0.1) for the decoder, and the composite pseudo-likelihood for the Potts head, which supervises the joint assignment at each directed edge against all *V* ^2^ candidate endpoint pairs. No token- or class-specific weights were applied, so protonation-state tokens occurred at their empirical frequencies in the training annotations.

### 4.3 Validation

#### Protonation-state recovery against neutron structures

Reference protonation states were assigned from deposited H/D atoms in neutron and joint X-ray/neutron structures. Histidine with a proton on both ring nitrogens was encoded as HIS-P and with exactly one as HIS-S; histidines with neither were excluded. Aspartate and glutamate with a proton on either carboxyl oxygen were encoded as ASP-P or GLU-P, and otherwise as ASP-D or GLU-D. A residue was excluded if no H/D atom was modeled anywhere in that residue. The resulting set comprised 216 structures and 7,434 labeled residues: 1,220 histidines, of which 416 were protonated imidazolium (34.1%); 3,200 aspartates, of which 36 were protonated (1.1%); and 3,014 glutamates, of which 58 were protonated (1.9%). Of these, 7,434 were present in the featurised structures and were scored.

Neutron entries were excluded from model training by the experimental-method filter and by a sequence filter removing structures overlapping the PDB training data. The protonation-state classifiers were, however, fitted on these structures, so classifier-derived labels for their cluster mates were produced by a classifier that had seen the benchmark. This evaluation therefore measures the transfer of that supervision into the sequence-design model on familiar folds, rather than independent validation of the labeling pipeline. Decoder probabilities were used directly; Potts probabilities were obtained by softmax over the negative conditional energy of Equation (2), with every other residue fixed at its experimentally derived reference token. Because only 1-2% of acidic residues were protonated, acid recovery was interpreted alongside the ranking benchmark below.

#### p*K*_a_ ranking benchmark

We assessed whether a fixed trained model ranked residues by experimental p*K*_a_ within each residue type, orienting all scores so that larger values indicate greater propensity for protonation. Experimental values were taken from PKAD [21]; of 1,024 measurements, 567 were primary measurements for the three residue types, and restriction to residues present in the parsed structures and assigned a value by PROPKA3 gave a common evaluation set of 101 histidines, 184 aspartates and 209 glutamates from 106 structures. Predictions were joined by PDB entry, chain, residue number and residue type, and structures overlapping the training data were removed.

Each structure was evaluated in a conditional-minus-self pass with the query residue’s token with-held from the context, and the p*K*_a_ proxy was the two-state energy gap between the protonated token *P* and its comparison token *C*,

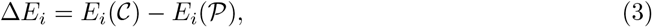

which is positive when the protonated state has the lower energy and is the log-odds of the protonated state under the two-state restriction. A tokens and canonical parent tokens were excluded. Three Potts scores were compared: the absolute protonated-state score *−E_i_*(*P*); the full gap; and an electrostatic gap omitting the field term and restricting both coupling sums to aspartate, glutamate, lysine, arginine and histidine partners. As a baseline, PROPKA 3.5.1 was run on the first model of each structure after re-serialization, so that all methods used the same coordinates.

#### Direct-coupling analysis

For each protein containing at least one titratable residue, an unpaired monomer alignment was generated with the ColabFold MMseqs2 API [35] and an evolutionary Potts model fitted by symmetric pseudo-likelihood maximization with *ℓ*_2_ regularization (*λ_h_* = 0.01, *λ_J_*= 0.02). Sequences were weighted by the reciprocal of the number of alignment members within 20% divergence, giving an effective count *M*_eff_. Couplings were symmetrized during fitting and each block placed in the zero-sum gauge, making couplings comparable across positions and families. Of 1,087 alignments fitted, 1,070 had *M*_eff_ *≥* 100 and were retained (median *M*_eff_ = 1,802). Matched structural and evolutionary coupling vectors were extracted over the 20 canonical partner types at each titratable site, sign-aligned, standardized per model and averaged to give a 20-value protonated-minus-comparison contrast for each; agreement was quantified by the Pearson correlation between the two contrasts.

#### Binder discrimination by Potts energy

We asked whether Potts energy distinguished experimentally confirmed binders from non-binders among designs the model did not help generate. The benchmark comprised 4,060 designed binders with experimental labels in four classes of differing prevalence: a meta-analysis set spanning 15 targets (3,486 designs; 11.0% binders), BindCraft designs (152; 42.8%), BoltzGen mini-binders (227; 29.1%) and BoltzGen nanobodies (195; 19.0%). Complexes were represented by Boltz-2 predicted structures under standard inference settings, and the ipTM from the same predictions served as the baseline, so that the two scores differ in scoring rather than input. Binder propensity was the negative Potts Hamiltonian of the complex normalized by its number of residues,

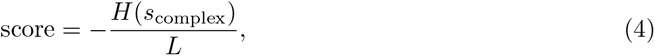

with larger values indicating a more favorable complex. Designs were pooled within each class and AP calculated against the experimental labels with a 2,000-sample bootstrap confidence interval.

### 4.4 Designing pH-sensitive binders with Proton-PottsMPNN

#### Design objective

A pH switch was formulated as a two-state design problem. One or more binder positions were fixed to target protonation-state tokens and the surrounding sequence redesigned to make the target assignment energetically preferable to its comparison assignment. Because the encoder is sequence-blind, each backbone was encoded once and its energy tables shared by all designs on that backbone, permitting exhaustive evaluation of alternative assignments. No coordinate noise was added during design.

For a fixed position *i* with target token *t_i_*and comparison set *C_i_*, the selective energy to be minimized was

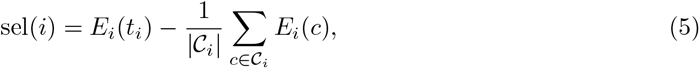

in which any token-independent contribution to the energy of position *i* cancels. Selectivity and stability were combined as

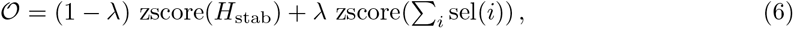

where *H*_stab_ is the Hamiltonian restricted to positions being changed and the two means and standard deviations were computed once from the joint block assignments and held fixed during the search. *λ* = 0 gives stability alone and *λ* = 1 selectivity alone. The switch direction is set entirely by the target and comparison tokens: targeting HIS-P relative to HIS-S favors a binder stabilized by histidine protonation, and the reverse assignment favors one destabilized by it. A dispersal penalty discouraging clustered charged residues was added where stated (Supp. Methods).

#### Placement

Placement selected the fixed positions and their target tokens, optionally restricted to a structural region of the binder chain. Interface positions were defined by C*α* atoms within 6 Å of a target-chain C*α* atom; core and surface positions were separated by relative solvent accessibility, with values below 0.2 defining the core. Candidates were ranked by one of two criteria. The Potts scan ranked each position by the selective-energy gap obtained by fixing it, evaluated against the current sequence. The decoder scan used the corresponding gap between decoder token log-probabilities after every designable binder position had been masked to the unknown token, making the ranking depend on the backbone and fixed target rather than on the seed sequence. For multiple centers, top-ranked candidates for each token were pooled and distinct-position combinations enumerated or sampled, each scored with all centers fixed simultaneously. Each retained plan defined the positions permitted to change, being the union of the centers’ coupled neighborhoods in both graph directions excluding the centers themselves, optionally restricted to the *k* nearest coupled neighbors per direction to control mutation load.

#### Sequence optimization

Two solvers optimized the permitted positions while holding all centers fixed at their target tokens. A third, hybrid solver is described in Supp. Methods.

##### MCMC Block descent

The default solver used block coordinate descent on Equation (6). Each permitted position was grouped with its most strongly coupled permitted partners in a block of size *B*, and the complete objective evaluated for all *V ^B^* joint assignments to the block. The minimizing assignment was accepted, or at nonzero temperature a Boltzmann sample over the joint assignments. Blocks were swept until a complete sweep produced no change or the maximum number of sweeps was reached.

##### Autoregressive infill

The autoregressive solver used the decoder as both sequence model and selectivity source. Centers were fixed at their target tokens, permitted positions set to the unknown token, and positions then filled in random order, each selected token becoming context for the next. Let ***t*** and ***c*** denote the simultaneous target and comparison assignments for all centers. For candidate token *a* at position *q* the decoder reward was 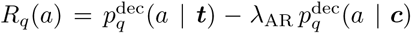, with both distributions evaluated under otherwise identical contexts, and the token sampled from a softmax over *R_q_*at temperature *T*. At *λ*_AR_ = 0 the token ranking matches the decoder probabilities, serving as the internal null; increasing *λ*_AR_ penalizes tokens favored under both assignments and enriches those specific to the target. No dispersal penalty was used with this solver, as the decoder’s learned sequence preferences discouraged low-complexity solutions.

##### Hybrid autoregressive infill

The hybrid variant uses the same masking and decoding order as autoregressive infill, but obtains its selectivity signal from the Potts head. At each designable position *q*, the decoder assigns a probability *p*_dec_(*a*) to each allowed token *a*, conditional on the pinned centers and positions already filled. The Potts head evaluates the selective energy at each pinned center after placing *a* at *q*. We combine these terms in the cost

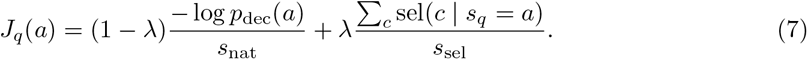

Here, sel(*c | s_q_* = *a*) is the target-versus-alternative energy gap of Equation (5) at pinned center *c*, evaluated with *a* at *q* and the other positions held at their current assignments. Each designable position lies in the neighborhood of at least one pinned center, so its token can change that center’s gap. The scales *s*_nat_ and *s*_sel_ are the standard deviations of the changes in decoder log-likelihoods and summed selective energy induced by single-position substitutions. They are computed once from the seed sequence with the centers pinned, before masking, and held fixed throughout decoding.

The next token is sampled from the allowed tokens *A_q_* according to

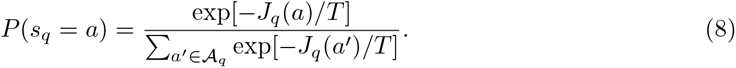

Thus, for finite *T >* 0, infilling is stochastic rather than a deterministic choice of the lowest-cost token.

### 4.5 Display of binder libraries

#### Yeast display library generation

All designs were synthesized as a multiplexed gene-fragment library at Twist Bioscience. Coding sequences were (GGS)*_n_*-padded to a uniform 360 nt and flanked by motifs matching the plasmid backbone to mediate library generation by homologous recombination. The reconstituted library was PCR-amplified over 15 cycles using primers annealing to the flanking regions. A Novo Nordisk-proprietary centromeric shuttle vector carrying a TRP1 auxotrophy marker was used for display, with designs fused to the N-terminus of the Aga2p surface anchor via a flexible linker under a galactose-inducible promoter: *Saccharomyces cerevisiae* EBY100 was grown to OD_600_ 1.2-1.6 in YPD at 30 *^◦^*C, washed twice in cooled water and once in electroporation buffer (1 M sorbitol, 1 mM CaCl_2_), conditioned for 30 min in 100 mM LiAc, 10 mM DTT, and washed twice more. Each reaction combined 400 µL of cells with 3 µg library DNA and 3.5 µg linearised backbone plasmid. Cells were recovered for 1 h in 1:1 1 M sorbitol/YPD, transferred to SD-CAA and cultured for 2 days.

#### Yeast library selections

Before each cycle, cells were adjusted to OD 0.5 in SG-CAA and induced overnight at 30 *^◦^*C. Three million induced cells were washed twice in PBS-A (PBS, 0.5% BSA) and incubated for 10 min at room temperature with 200 nM biotinylated PD-L1 (AcroBio, PD1-H82E5). Cells were stained with either Streptavidin-PE (BD Pharmingen, 554061) and AF647 anti-DYKDDDDK (BioLegend, 637316), or Streptavidin-AF647 (Invitrogen, 10349222) and AF488 anti-DYKDDDDK (BioLegend, 637318), to visualize target binding and surface expression, all at 1:200 in PBS-A. For enrichment of pH-sensitive binders, all washing and incubation steps used PBS-A at pH 5.0; for pH binning, the same steps were performed at a defined pH between 4.0 and 7.4. For isolation under physiological conditions, all steps used PBS-A at pH 7.4, with 1000 nM biotinylated PD-L1 tetramerised beforehand with 250 nM streptavidin-PE or streptavidin-AF647. Sorting used a Sony MA900 cell sorter, and isolated cells were recovered for 48 h in SD-CAA.

#### MACS library selections

All incubation, washing and capture steps used PBS-A at pH 5.0. Twenty million induced cells were washed twice in PBS-A and incubated for 10 min at room temperature with 1000 nM biotinylated PD-L1, then re-suspended in 900 µL PBS-A with 100 µL Streptavidin MicroBeads (Miltenyi, 130-048-102) for 15 min at 4 *^◦^*C. Cells were applied to an LS column (Miltenyi, 130-042-401) on a QuadroMACS separator, and non-specifically captured cells depleted with 8 mL PBS-A. Cells expressing a pH-responsive design were eluted with 2*×*5 mL PBS-A at pH 7.4. All sorting pools were sequenced on an Illumina MiSeq using the Reagent Kit v3 (600-cycle).

#### NGS data processing

Sorted pools were sequenced in two campaigns and processed separately throughout, since sample indices collide between them and denote different selections. The first comprised the initial library, an expression-only sort and the FACS binding and pH-binning sorts, split across two MiSeq runs; the second comprised the MACS rounds and a pH 7.4 track. Counts were never pooled across campaigns. Libraries were sequenced on a MiSeq (Reagent Kit v3, 600 cycles) in a 300 + 8 + 8 + 300 configuration with unique dual 8 nt indices. Base calling and demultiplexing were performed on-instrument with Illumina Local Run Manager (GenerateFASTQ v3.1.0) in FASTQ-only mode, assigning reads by exact-to-one-mismatch matching of both indices. Adapter trimming against the Nextera mosaic end removed *<*0.1 % of bases, as the 360 nt amplicon exceeds the paired read length. Demultiplexing was audited by counting read pairs in every index well and characterising the undetermined pool. The two first-campaign runs proved complementary rather than replicate, each sample carrying 0.3 to 2.5 M reads in one run and 0 to 155 index-bleed reads in the other, so the run used for each sample was selected by depth.

Because the amplicon has constant flanks and a fixed 360 nt insert, read pairs were merged by exact positional stitching rather than overlap discovery. R1 was anchored on the constant sequence immediately 5*′* of the insert and the reverse complement of R2 on the vector (G_4_S) linker immediately 3*′* of it, with the FLAG tag as a fallback anchor. Where the *∼*160 nt doubly covered region disagreed, the higher Phred base was taken. Read pairs lacking either anchor, or not jointly spanning the insert, were discarded.

Per-design counts were taken from exact protein matches only. Translations that were not exact library members were assigned to their nearest member by Hamming distance up to 6 substitutions, considering only distinct translations observed at least twice. This rescue was used to define the assigned-read denominator and to quantify what fraction of unassigned reads were error-laden reads of genuine members, but was not folded into per-design counts. All cross-sample comparisons used counts per million assigned reads. Designs were called present against a per-design background model in the spirit of EmptyDrops [36], rather than a flat read floor, since stray reads from index hopping scale with a design’s abundance in co-loaded samples rather than with the sample itself. Consistent with this, low counts tracked neighbor abundance (Spearman 0.32 to 0.63) and not input-library abundance. For design *d* in sample *s* the expected stray count is *λ_ds_* = *h∑_j_ N_j_f_dj_*, over donor samples *j* with depth *N_j_* and observed frequency *f_dj_*, where *h* is the hopping rate. Under the null that the design is absent the observed count is Poisson with this rate, giving *p* = *P* (Poisson(*λ_ds_*) *≥ n_ds_*). These were corrected across designs within a sample by Benjamini–Hochberg and a design called present at a false-discovery rate of 0.01. Calls were repeated at *h* ∈ {0.5, 1, 2} and only conclusions stable across that range retained, with *λ_ds_* floored at 0.05.

## Acknowledgments

We thank Green Ahn for generously sharing structural data for pH-sensitive binders, which contributed to this work.

## Code and model availability

Proton-PottsMPNN is available at https://github.com/christian-creator/ProtonPottsMPNN under the MIT license, together with the trained weights and the histidine and acidic-residue protonation-state classifiers used to generate the training annotations.

## Author contributions

C.P.J. trained and benchmarked the models and designed the PD-L1 binders. L.D. performed the yeast-display FACS screening and the quality-control experiments. U.J. performed the MACS screening. L.D. and U.J. performed the next-generation sequencing. F.B. helped conceptualize the analysis. T.P.J. and S.O. supervised the project. C.P.J. wrote the original draft of the manuscript. T.F and L.G is currently testing pH-sensitive binders. All authors reviewed and approved the final manuscript.

## 5 Supplementary

**Table 1.**
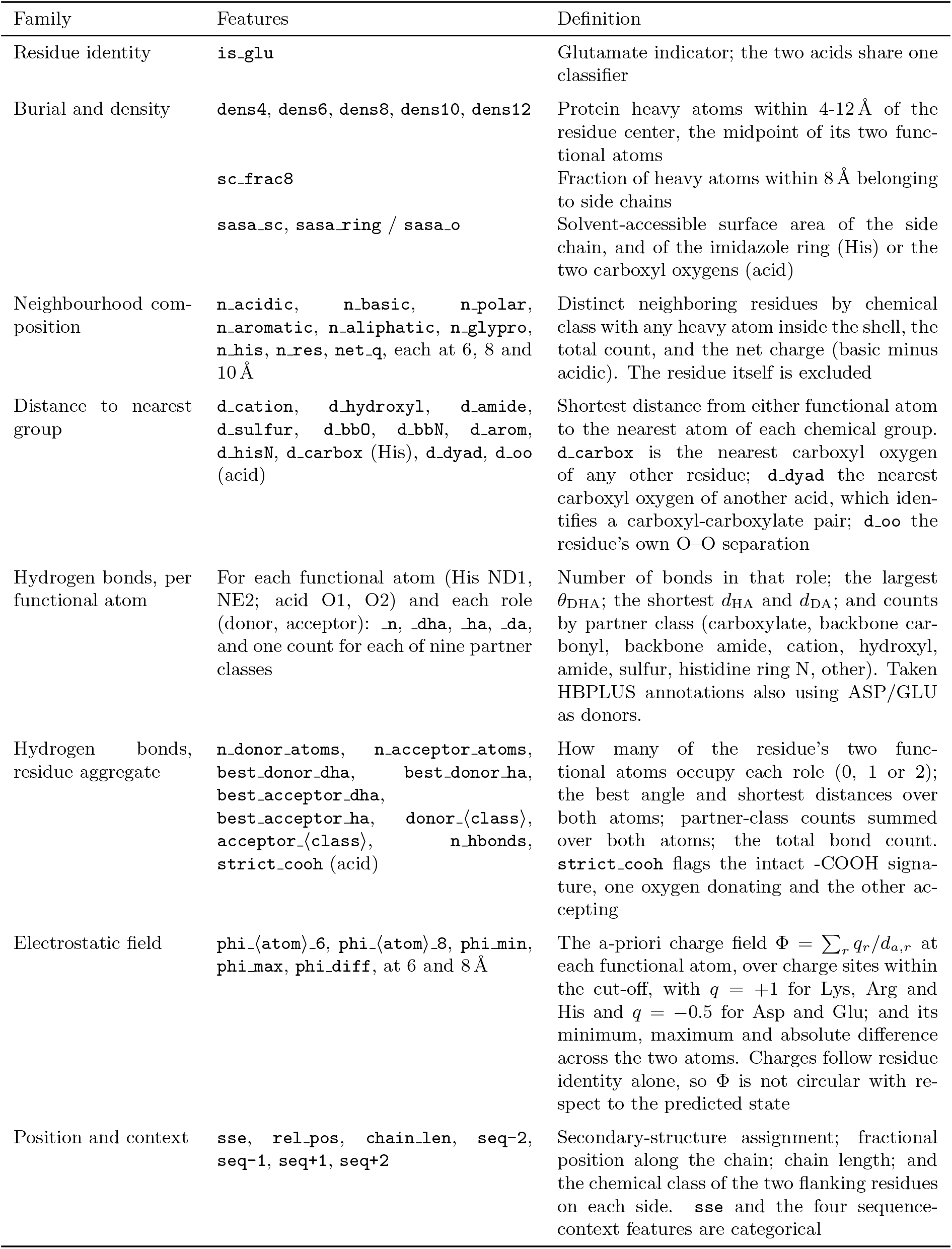
Geometric features for Protonation predictor. Feature manifest for the protonation-state classifiers. All features are computed from protein heavy atoms of the twenty canonical amino acids; hydrogens are removed beforehand, and waters, ligands, cofactors and metals never enter a feature. Functional atoms are the two ring nitrogens ND1 and NE2 for histidine and the two carboxyl oxygens for aspartate and glutamate. Hydrogen bonds are enumerated at the permissive criteria of the annotation pool (*d*_HA_ *≤* 3.2 Å, *d*_DA_ *≤* 4.0 Å, *θ*_DHA_ *≥* 90*^◦^*) and passed to the classifier unfiltered.

**Fig. S1.**
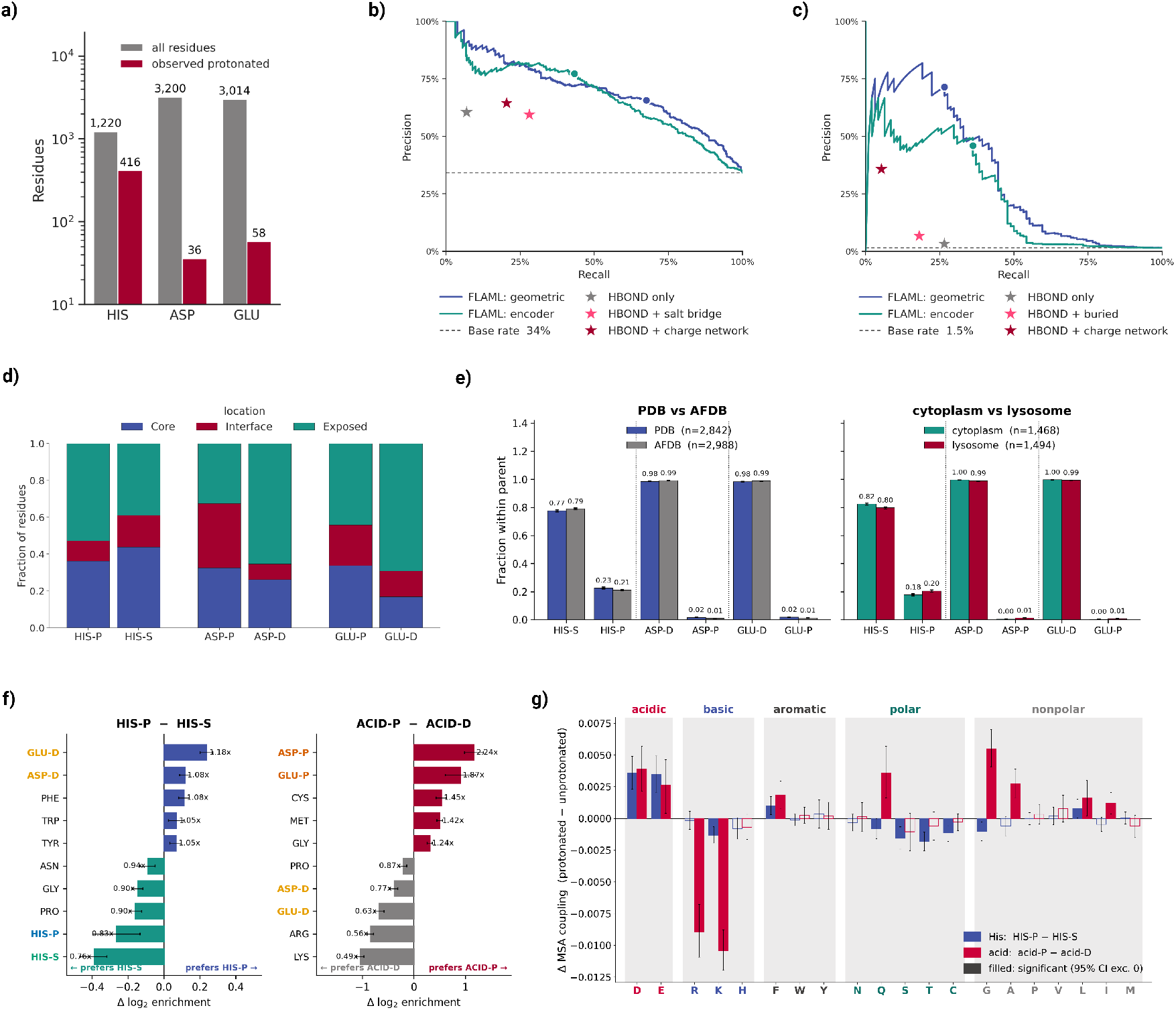
Evaluation of protonation-state labels. **a)** Numbers of histidine (HIS), aspartate (ASP), and glutamate (GLU) residues in the experimental H/D-resolved dataset (grey) and observed in the protonated state (orange). **b,c)** Out-of-fold precision-recall curves from grouped five-fold cross-validation for histidine (**b**) and pooled ASP/GLU _(_**_c_**_) classifiers using geometric features (blue) or ProteinMPNN encoder embeddings (orange). Stars denote the rule-based methods indicated in each key, dashed lines show protonated-state prevalence, and circles mark the selected thresholds._ **_d)_** _Fractions of experimentally labeled residues in core (blue), interface (orange), and exposed (green) locations; sample sizes are shown below the bars._ **_e)_** _Fractions assigned to each protonation state in PDB versus AFDB structures (left) and cytoplasmic versus lysosomal AFDB proteins (right). Error bars show bootstrap standard deviations; sample sizes are given in the keys._ **_f)_** _Differences in log2 neighborhood enrichment between HIS-P and HIS-S, and between pooled protonated and deprotonated acidic residues (ACID-P and ACID-D). Error bars are confidence intervals from 500 paired bootstrap resamples._ **_g)_** _MSA-derived coupling differences between protonated and de-protonated states for histidine (blue) and acidic residues (red). Error bars are 95% bootstrap confidence intervals; filled bars indicate intervals excluding zero._

**Fig. S2.**
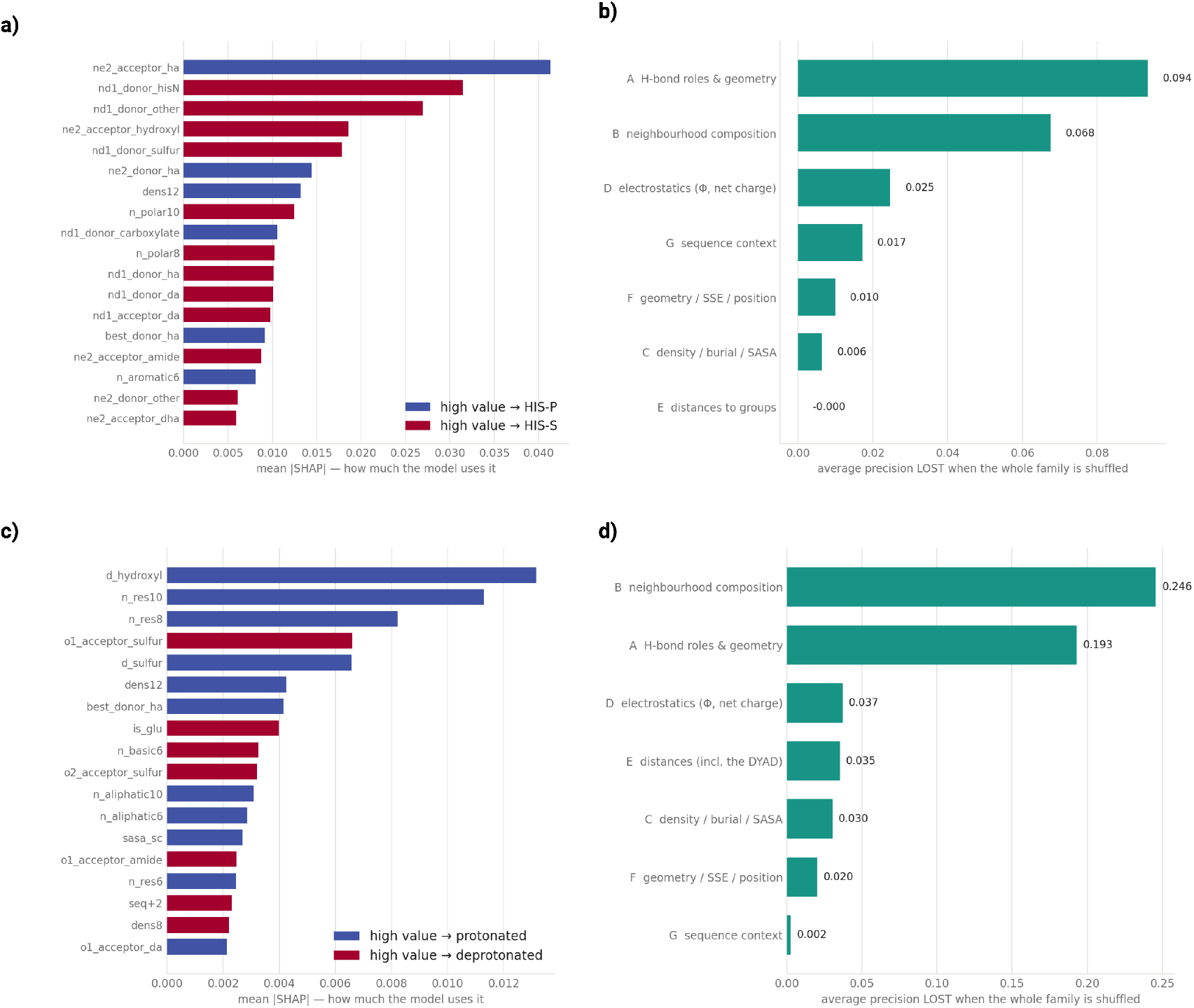
FLAML shap analysis: **a,c)** The twenty features with the highest mean |SHAP| for the histidine (**a**) and pooled aspartate/glutamate (**c**) classifiers, measuring how strongly the model uses each feature. Bars are colored by the direction of the association, with blue indicating that high values favor the protonated state and red the deprotonated state. **b,d)** Loss in average precision when each feature family is shuffled in turn, for histidine (**b**) and pooled aspartate/glutamate (**d**).

**Fig. S3.**
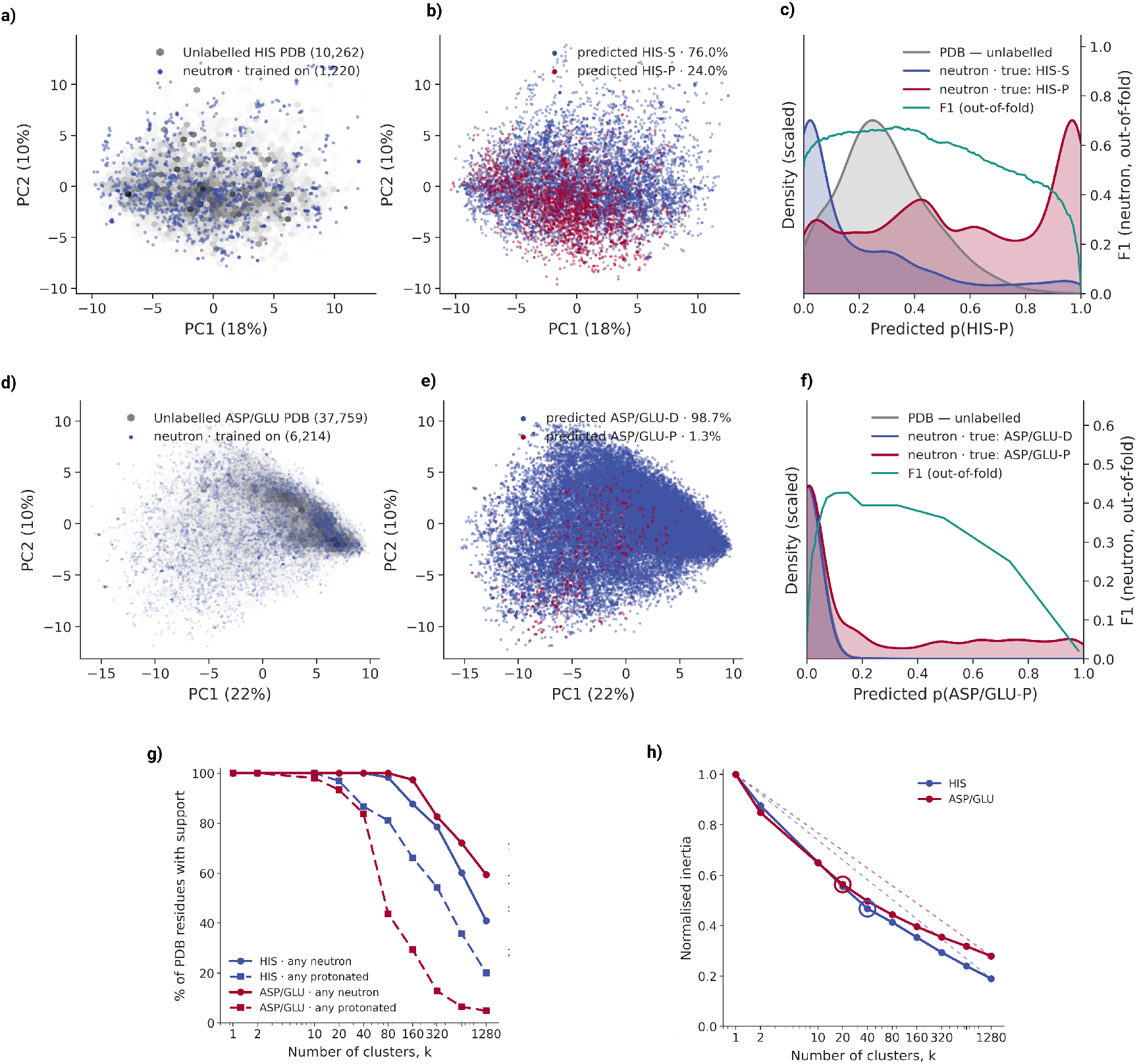
Coverage of the labeled residues within the wider PDB. **a,d)** Principal-component analysis of the geometric features used by the protonation-state classifiers, for histidine (**a**; 138 features) and pooled aspartate and glutamate (**d**; 141 features). Blue points are the labeled residues from neutron and joint X-ray/neutron structures on which the classifiers were trained (1,220 histidines; 6,214 acidic residues); grey density shows unlabeled residues sampled from the wider PDB (10,262 histidines; 37,759 acidic residues). **b,e)** The same projections for the unlabeled PDB residues, colored by the protonation state assigned by the XGBoost classifiers. **c,f)**Distributions of predicted protonation probability for histidine (**c**) and pooled acidic residues (**f**). Blue and red show out-of-fold predictions for labeled residues in the deprotonated and protonated states, respectively; grey shows the unlabeled PDB residues. Green gives the out-of-fold F1 score as a function of the assignment threshold (right axis). **g)** Fraction of PDB residues falling in a *k*-means cluster of the feature space that contains at least one labeled residue (solid) or at least one labeled protonated residue (dashed), as a function of the number of clusters. **h)** Normalized inertia against cluster number; circles mark the elbow.

**Fig. S4.**
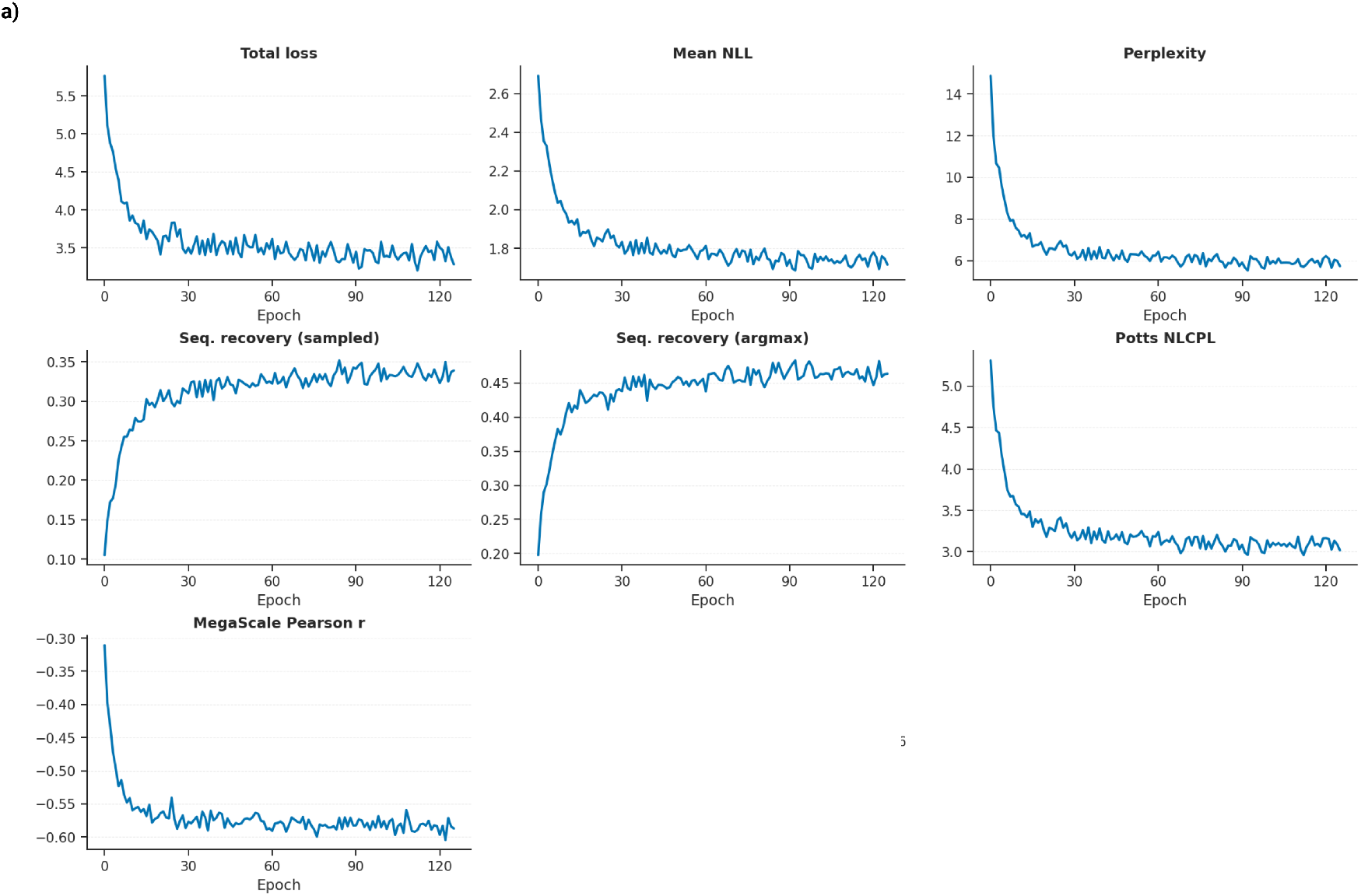
Training curves for Proton-PottsMPNN. Validation metrics over 125 training epochs: total loss; mean negative log-likelihood and perplexity of the autoregressive head; full-vocabulary sequence recovery under sampled and argmax decoding; the negative composite pseudo-log-likelihood of the Potts head; and the Pearson correlation between the Potts energy change and experimental ΔΔ*G* on the Megascale set.

**Fig. S5.**
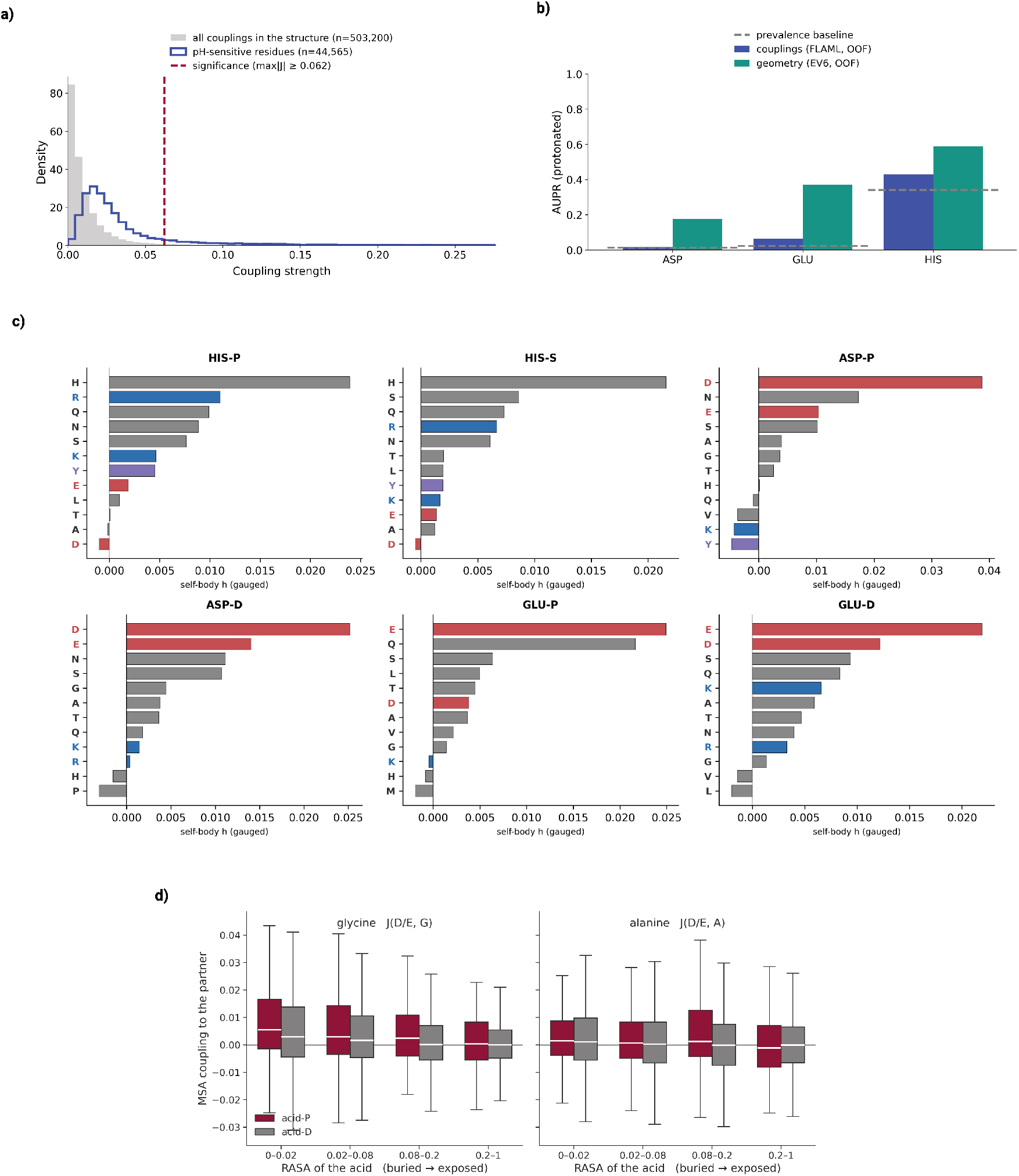
Evolutionary couplings at titratable positions. **a)** Distribution of evolutionary coupling strengths involving predicted protonated residues (blue; *n* = 44,565) against all couplings in the same structures (grey; *n* = 503,200). The dashed line marks the 95th percentile of the all-coupling distribution, used as a significance threshold. **b)** Average precision for protonation-state prediction by classifiers trained on evolutionary couplings (blue) against the geometry-based classifiers used to label the Proton-PottsMPNN training data (green), evaluated out-of-fold on residues with deposited protonation states. Dashed lines show protonated-state prevalence; *n* gives the number of residues and proteins per residue type. **c)** Gauged single-site preferences from the fitted DCA models, averaged over sites assigned to each protonation state. Partner residues are colored by charge, with acidic in red and basic in blue. **d)** Evolutionary coupling to glycine and to alanine as a function of the relative solvent accessibility of the acidic residue, separated by assigned protonation state.

**Fig. S6.**
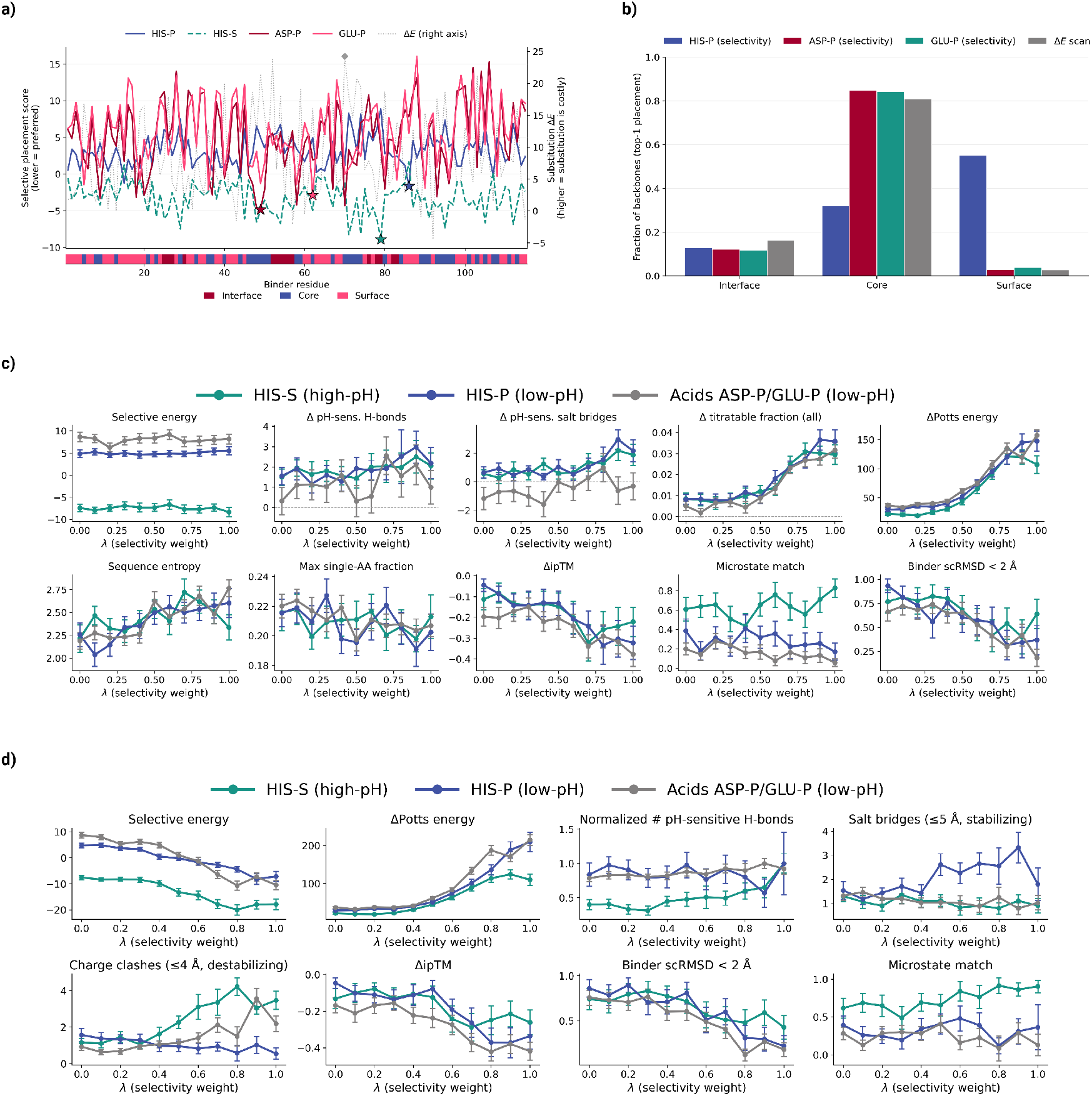
Placement of titratable centers and the effect of λ under autoregressive and hybrid redesign. **a)** Per-position placement scores along one example binder, ranked by the selective energy for HIS-P, HIS-S, ASP-P and GLU-P (left axis, lower is preferred) and by the state-agnostic substitution Δ*E* (right axis, higher is costly). Stars mark the top-ranked position for each criterion. The bar beneath the axis indicates whether each residue lies at the interface, in the core or on the surface. **b)** Fraction of top-ranked placements falling in each structural region, pooled across the 37 seed binders, for the three selectivity criteria and the Δ*E* scan. **c, d)** Effect of the selectivity weight *λ* on energetic, sequence and structural metrics for designs generated with the autoregressive decoder (c) and with hybrid autoregressive infilling, in which the decoder proposes tokens and the Potts head supplies the selectivity signal (d). The Potts block-descent counterpart is shown in Fig. **3**a. Teal, blue and grey denote HIS-S (high-pH target state), HIS-P and pooled ASP-P/GLU-P (low-pH) designs. Δ values are relative to the seed, error bars show the standard error across designs.

**Fig. S7.**
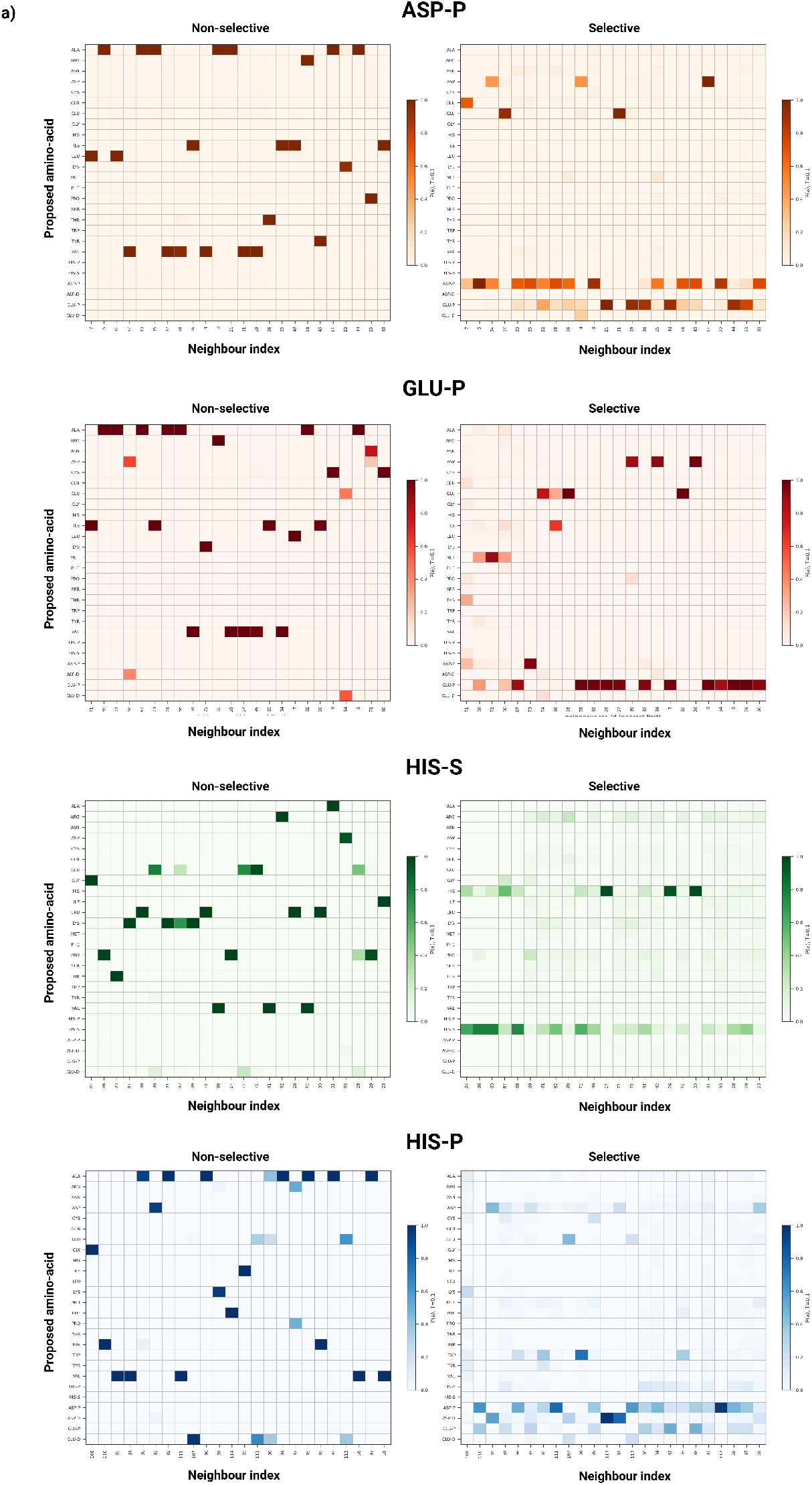
Amino-acid preferences around a pinned titratable center. Token preferences at neighboring positions when redesigning around a center fixed at ASP-P, GLU-P, HIS-S or HIS-P. For each center, the left panel shows preferences under the non-selective objective, which optimizes the global Potts energy alone (*λ* = 0), and the right panel under the selective objective, which optimizes the selective energy alone (*λ* = 1). Rows give the candidate token at the neighboring position and columns its graph-neighbor rank, ordered from nearest to furthest from the center; color gives the preference for that token at that position.

**Fig. S8.**
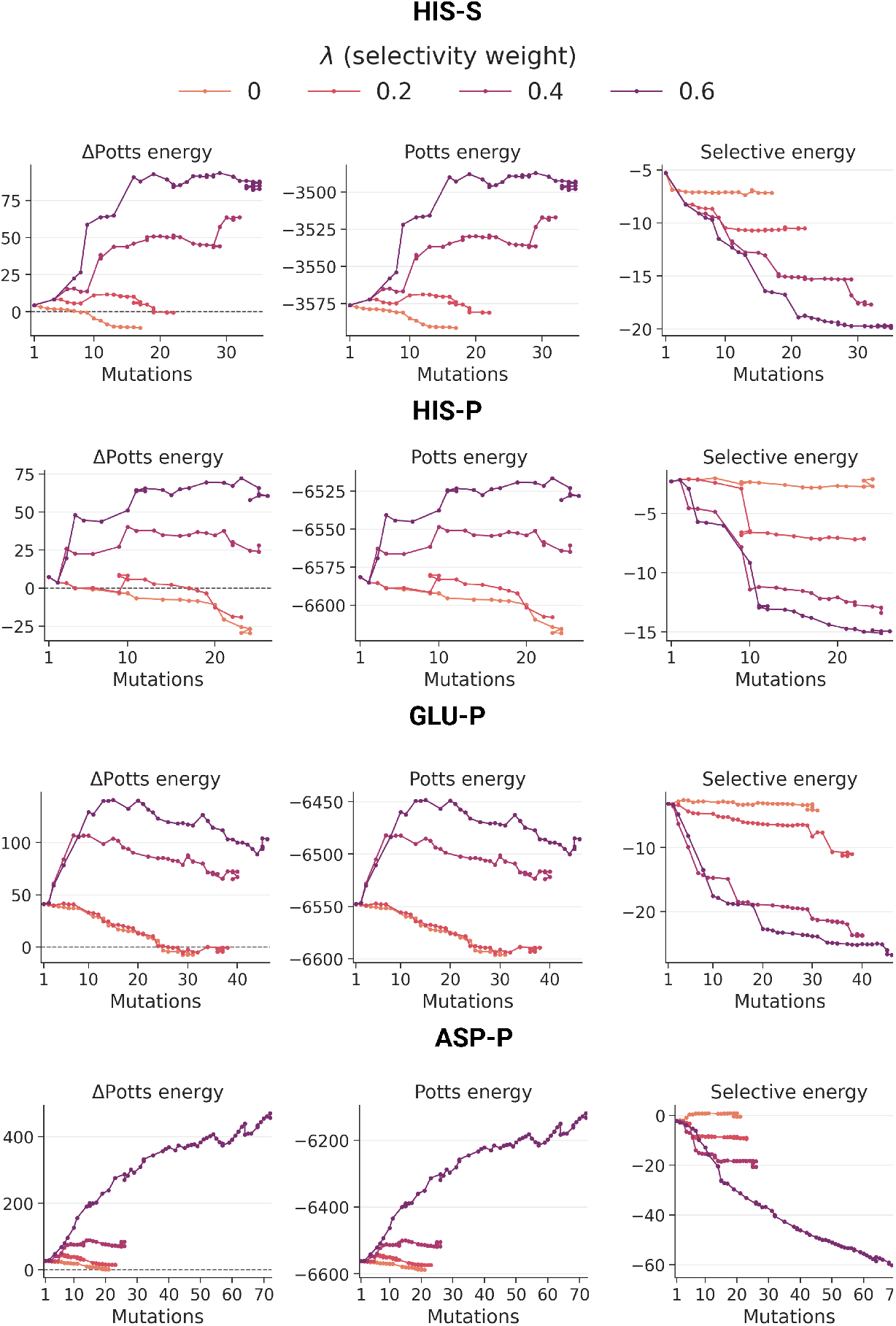
Example optimization trajectories. Block-descent trajectories for a example seed binders redesigned around a center pinned at HIS-S, HIS-P, GLU-P or ASP-P, at four values of the selectivity weight *λ*. For each target state, the change in Potts energy relative to the seed (left), the absolute Potts energy (center) and the selective energy of the pinned center (right) are shown against the number of accumulated mutations. Lower Potts energy and lower selective energy are more favorable; the dashed line marks the seed Potts energy. Trajectories end when a complete sweep produced no further change.

**Fig. S9.**
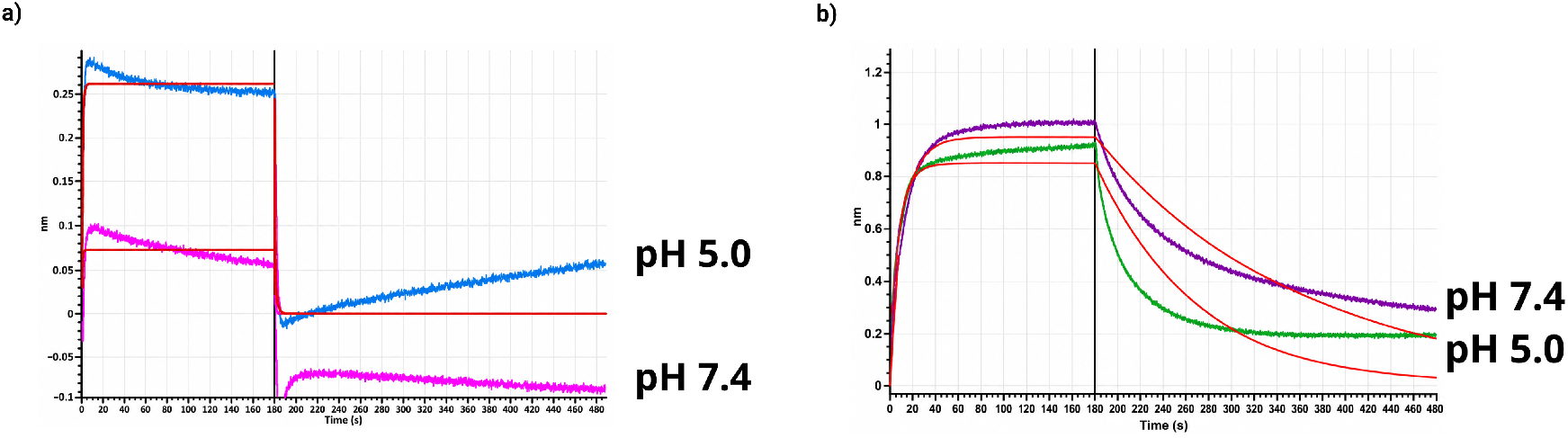
The PD-L1 reagent engages its native ligand at both selection pH values. Biolayer interferometry of biotinylated PD-L1 loaded onto streptavidin biosensors and dipped into recombinant PD-1 at pH 5.0 or pH 7.4. Association was measured for 180 s, to the vertical line, and dissociation for a further 300 s in ligand-free buffer; red lines are 1:1 fits. **a)** Monomeric His-tagged human PD-1 at 5 µM. **b)** Human PD-1 fused to mouse Fc IgG2a at 1 µM, which engages PD-L1 avidly.

**Fig. S10.**
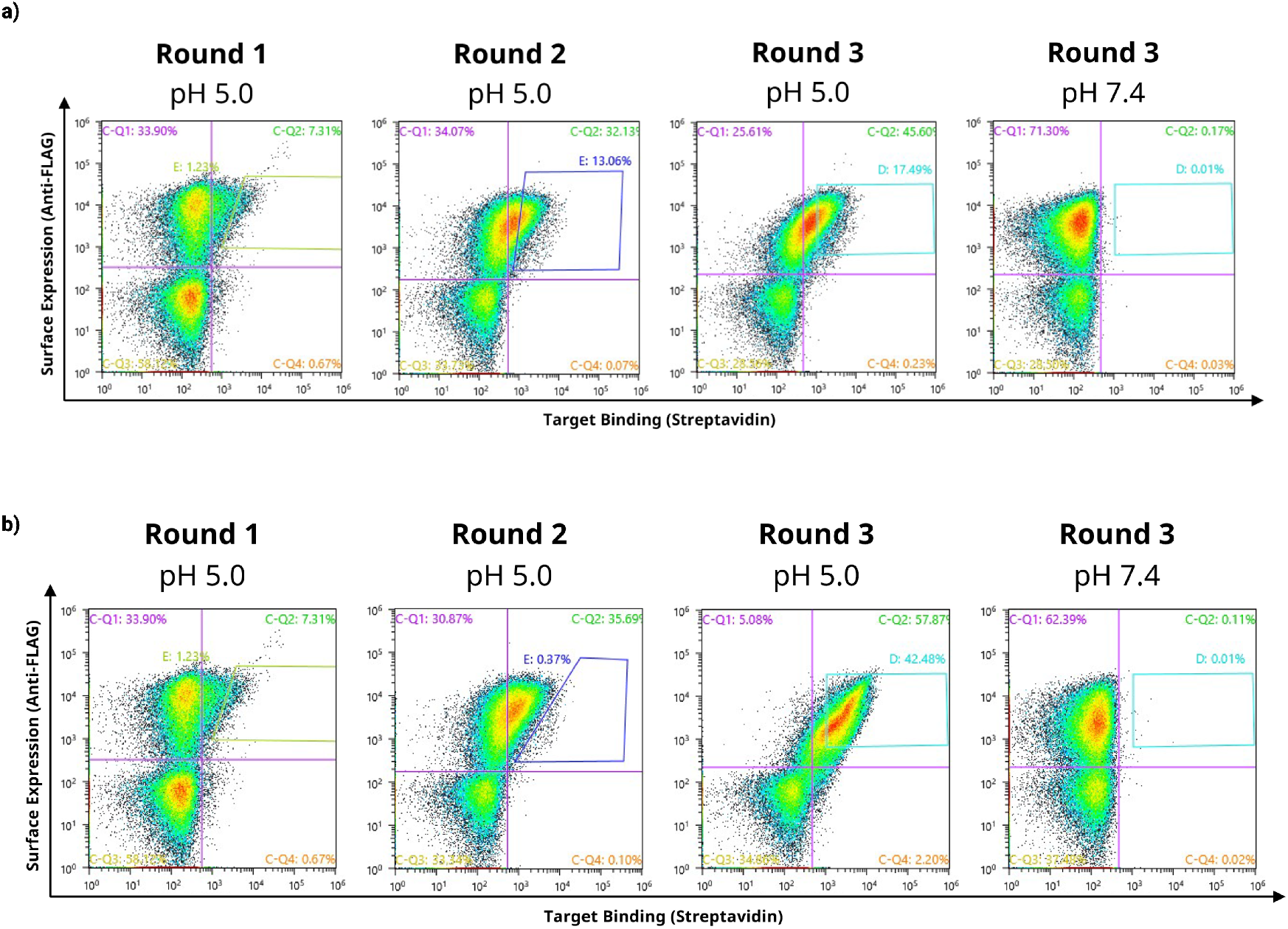
Selection of the PD-L1 library at pH 5.0. Flow cytometry of the yeast-display library over three consecutive rounds of sorting, following incubation with biotinylated PD-L1. Axes show target binding (streptavidin) and surface expression (anti-FLAG); percentages give the fraction of events in the indicated region, and the drawn gate in each round-2 panel is the population collected. **a)** Selection using an inclusive binding gate, which retains weak binders. **b)** Selection from the same round-1 population using a stringent gate, which retains only the highest-binding cells, collecting 0.37% of events against 13.06% under the inclusive gate. The rightmost panel in each row shows the enriched population assayed at pH 7.4, where binding is undetectable.

**Fig. S11.**
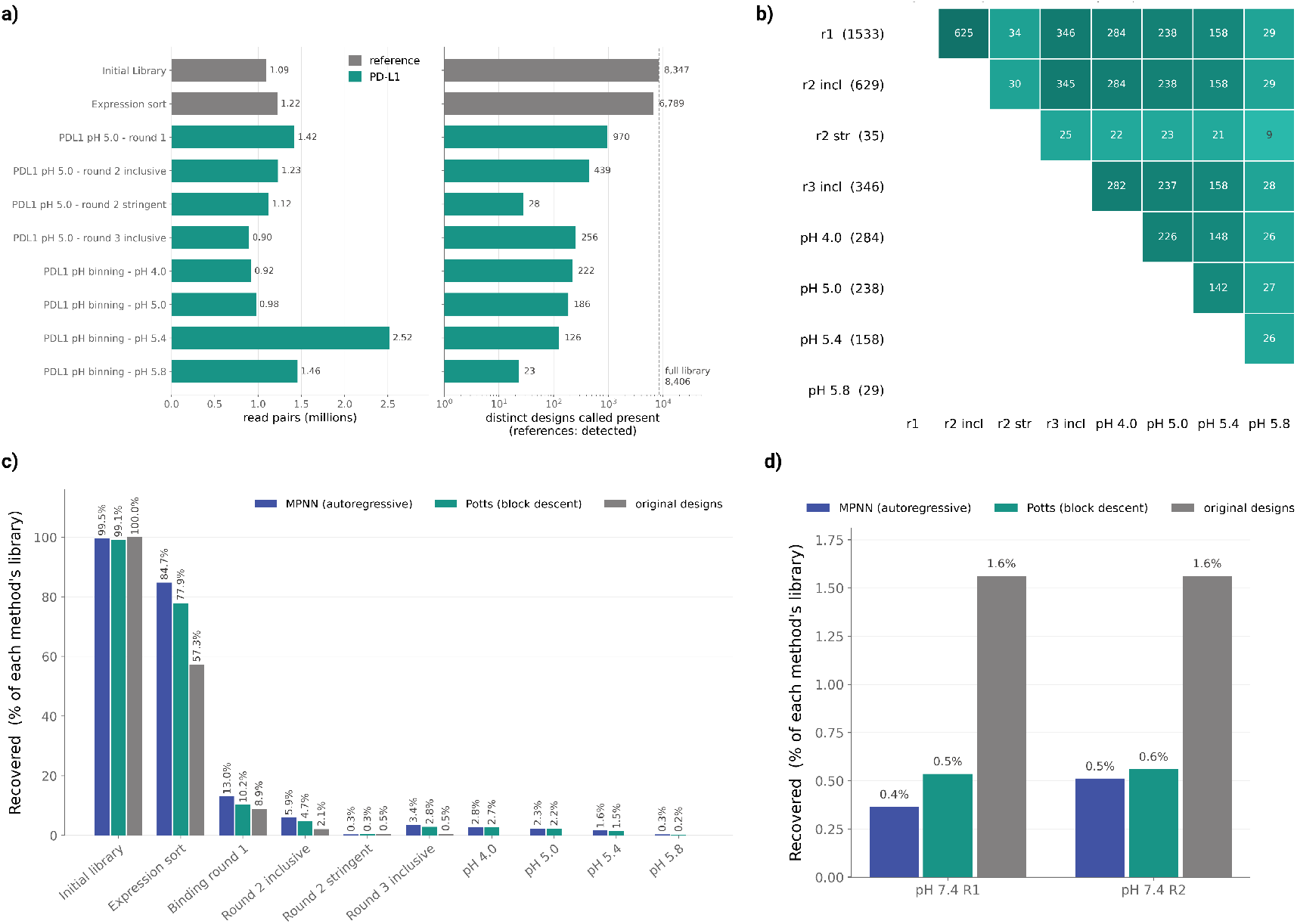
Sequencing of the PD-L1 sorting pools and recovery by design strategy. **a)** Sequencing summary for the initial library, the expression sort, the three binding rounds at pH 5.0 and the four pH bins. Left, read pairs obtained per pool. Right, the number of distinct sequences called present in each pool on a logarithmic scale, separated into PD-L1 designs (teal) and the reference sequences carried in the same synthesis (grey); the dashed line marks the 8,407 designs in the assembled library. **b)** Designs detected in both pools of a pair, for the three binding rounds (round 1; round 2 under the inclusive and the stringent gate; round 3 inclusive) and the four pH bins. Parentheses in the row labels give the number of distinct designs detected in each pool, and each cell the number shared with the pool in the corresponding column. **c)** Recovery by design strategy across the same pools, for designs redesigned with the hybrid autoregressive decoder (blue) or with Potts block descent (teal) and for the unmodified ProteinMPNN seed sequences included as controls (grey). Each bar gives the designs of that class detected in the pool as a percentage of the designs of that class present in the library, so that classes of different size are comparable. **_d)_** _The same quantity for the two rounds of selection carried out at pH 7.4 with 1,000 nM tetramerised PD-L1._

**Fig. S12.**
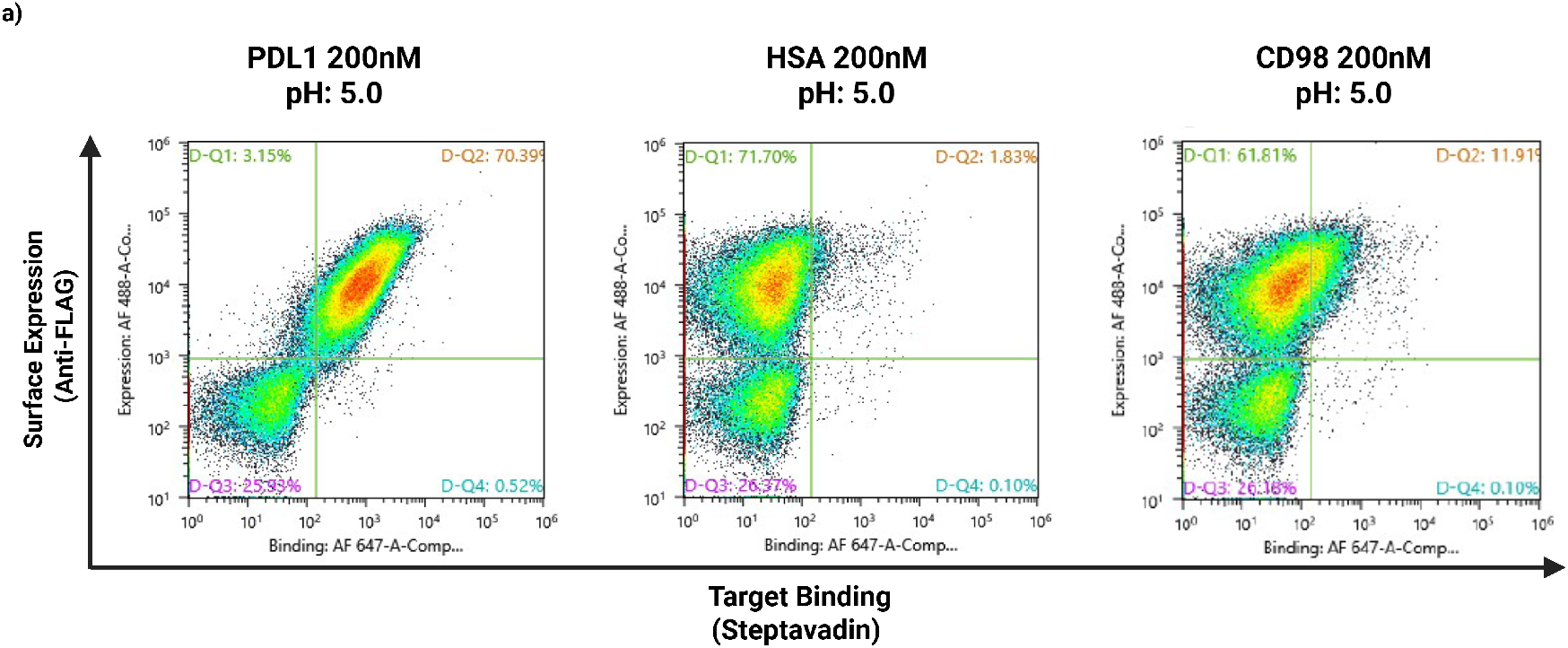
Binding of the enriched library to off-target proteins. Flow cytometry of the enriched yeast-display library following incubation with 200 nM biotinylated PD-L1, human serum albumin (HSA) or CD98, with all incubation and washing steps at pH 5.0. Axes show target binding (streptavidin–AF647) and surface expression (anti-FLAG–AF488).

**Fig. S13.**
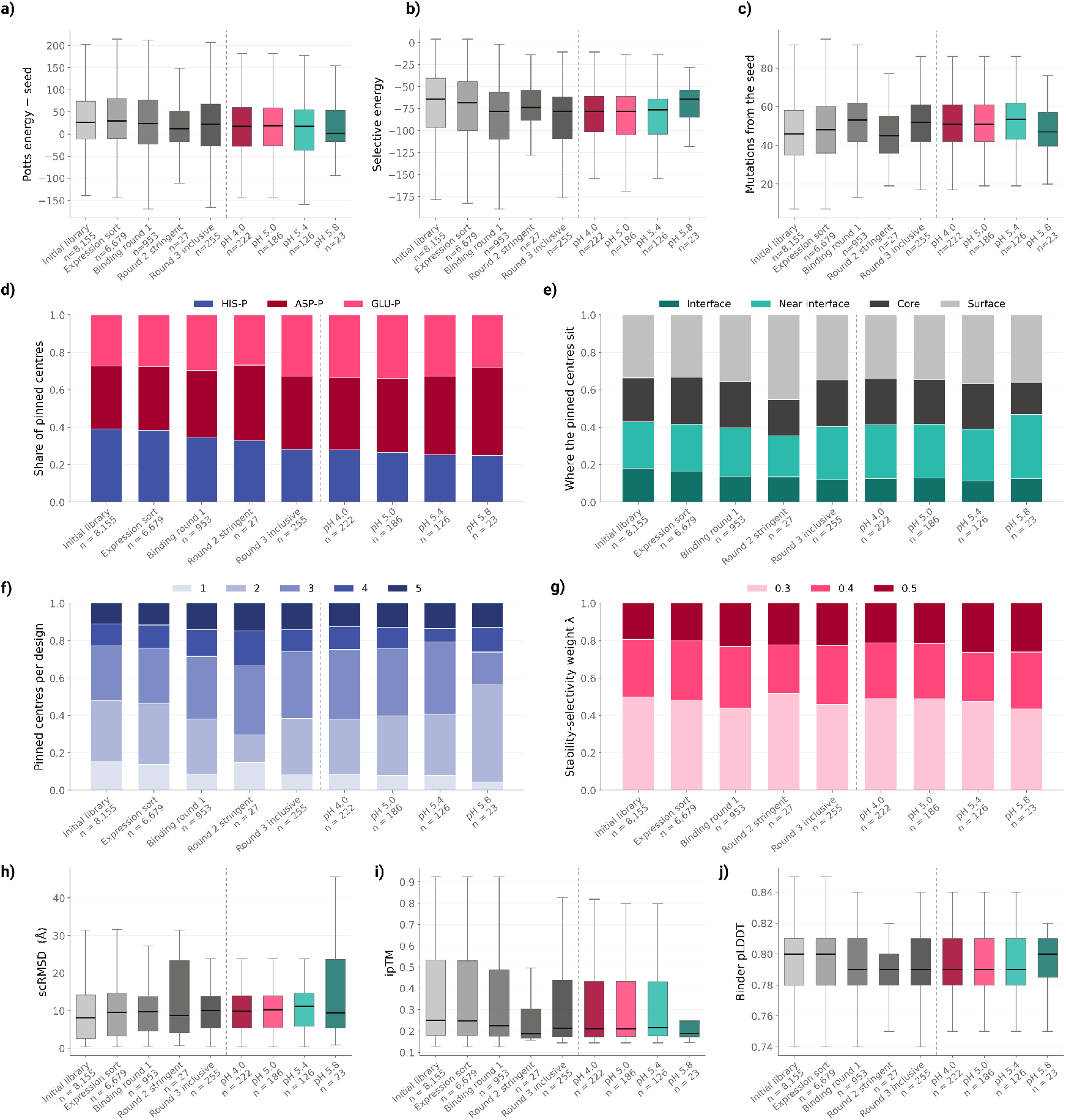
Properties of the PD-L1 library across sorting pools. **a–c)** Distributions of the change in Potts energy relative to the seed sequence (**a**), selective energy summed over the pinned centers (**b**) and number of mutations from the seed sequence (**c**), for the initial library, the expression sort, the three binding rounds and the four pH bins. The dashed line separates the binding rounds from the pH series, and *n* gives the number of designs detected in each pool. **d–g)** Composition of the same pools by the protonation state of the pinned centers (**d**), their structural location (**e**), the number of pinned centers per design (**f**) and the stability–selectivity weight *λ* used to generate them (**g**). **h–j)** Distributions of self-consistency RMSD (**h**), ipTM (**i**) and binder pLDDT (**j**) across the same pools. These structures were predicted after selection for analysis only; no structure-prediction filter was applied to the library at any stage.

**Fig. S14.**
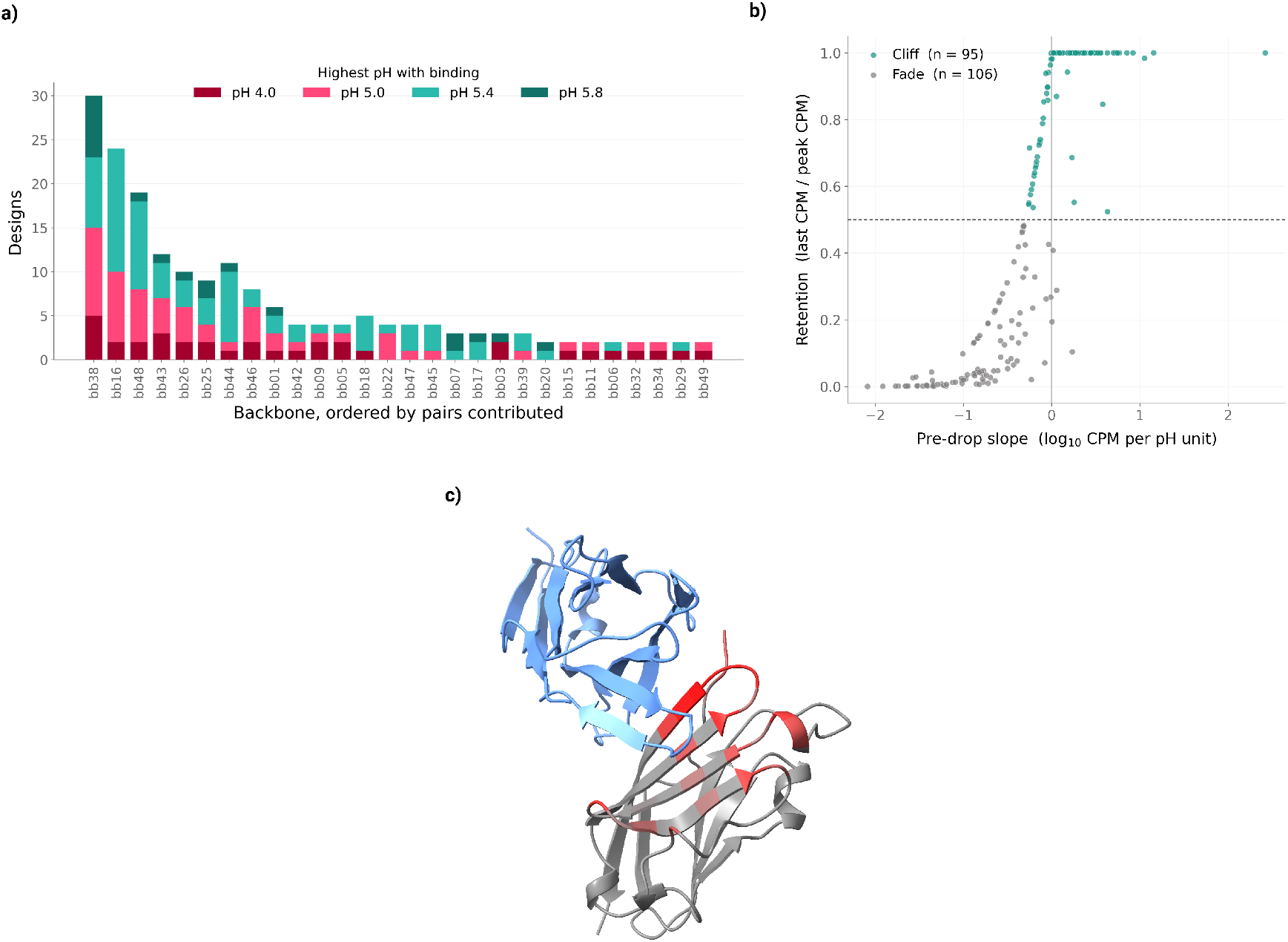
Backbone distribution and shape of the pH response. **a)** Number of designs recovered from each backbone contributing more than one sequence, ordered by contribution and colored by the highest pH at which binding was retained. **b)** Classification of the pH response by its shape. The vertical axis gives retention, the abundance in the last bin in which a design was detected divided by its highest abundance across bins, so that values near one indicate a design that was still at peak abundance when binding was lost. The horizontal axis gives the pre-drop slope, the change in log_10_ CPM per pH unit over the three bins preceding the loss, with more negative values indicating a gradual decline. Designs above the retention threshold are classified as cliffs (*n* = 95) and those below as fades (*n* = 106).**c)** Predicted structure of PD-L1 (grey) in complex with an example recovered binder (blue), with PD-L1 surface residues colored by how frequently they are contacted across the recovered designs.

**Fig. S15.**
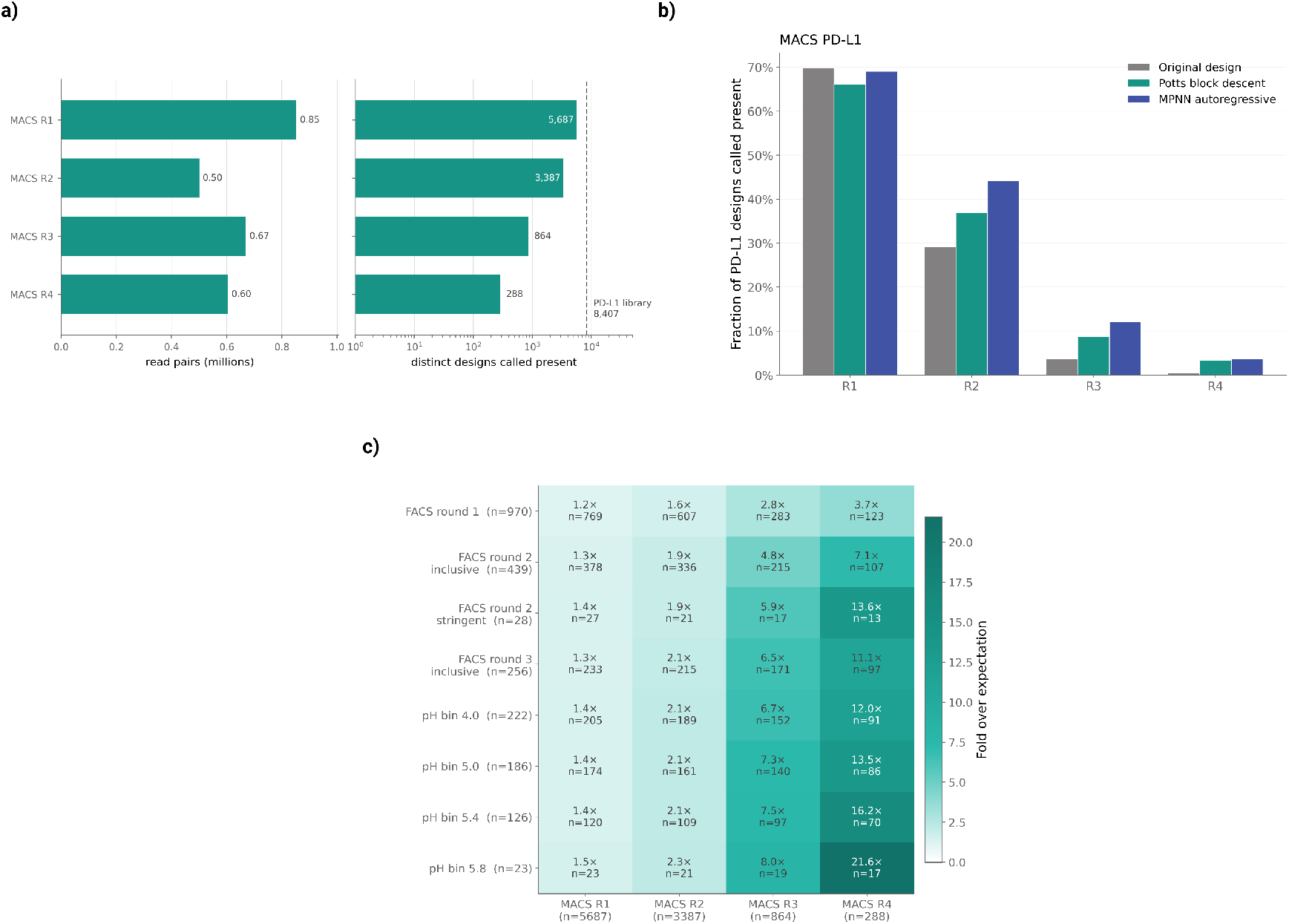
Magnetic selection of the PD-L1 library. **a)** Sequencing summary for the four rounds of magnetic-activated cell sorting. Left, read pairs obtained per round. Right, the number of distinct designs called present in each round on a logarithmic scale; the dashed line marks the 8,407 designs in the assembled library. **b)** Fraction of designs called present at each round, shown separately for the unmodified ProteinMPNN seed sequences included as controls (grey) and for designs redesigned with Potts block descent (teal) or with the autoregressive decoder hybrid approach (blue). Each bar gives the designs of that class detected in the round as a percentage of the designs of that class present in the library, so that classes of different size are comparable. **c)** Overlap between the fluorescence- and magnetic-sorting tracks. Rows are the FACS binding rounds and pH bins, columns the four MACS rounds, and *n* in each label gives the number of distinct designs detected in that pool. Each cell gives the number of designs detected in both pools and the fold by which this exceeds the overlap expected if the two pools drew independently from the library. **d)** Standardised difference in design properties between the round-4 MACS pool and the input library (dark blue) or the round-1 MACS pool (teal). Positive values indicate higher values in round 4. Points give the mean difference and horizontal bars its 95% confidence interval; properties are ordered by the difference against the library.

**Fig. S16.**
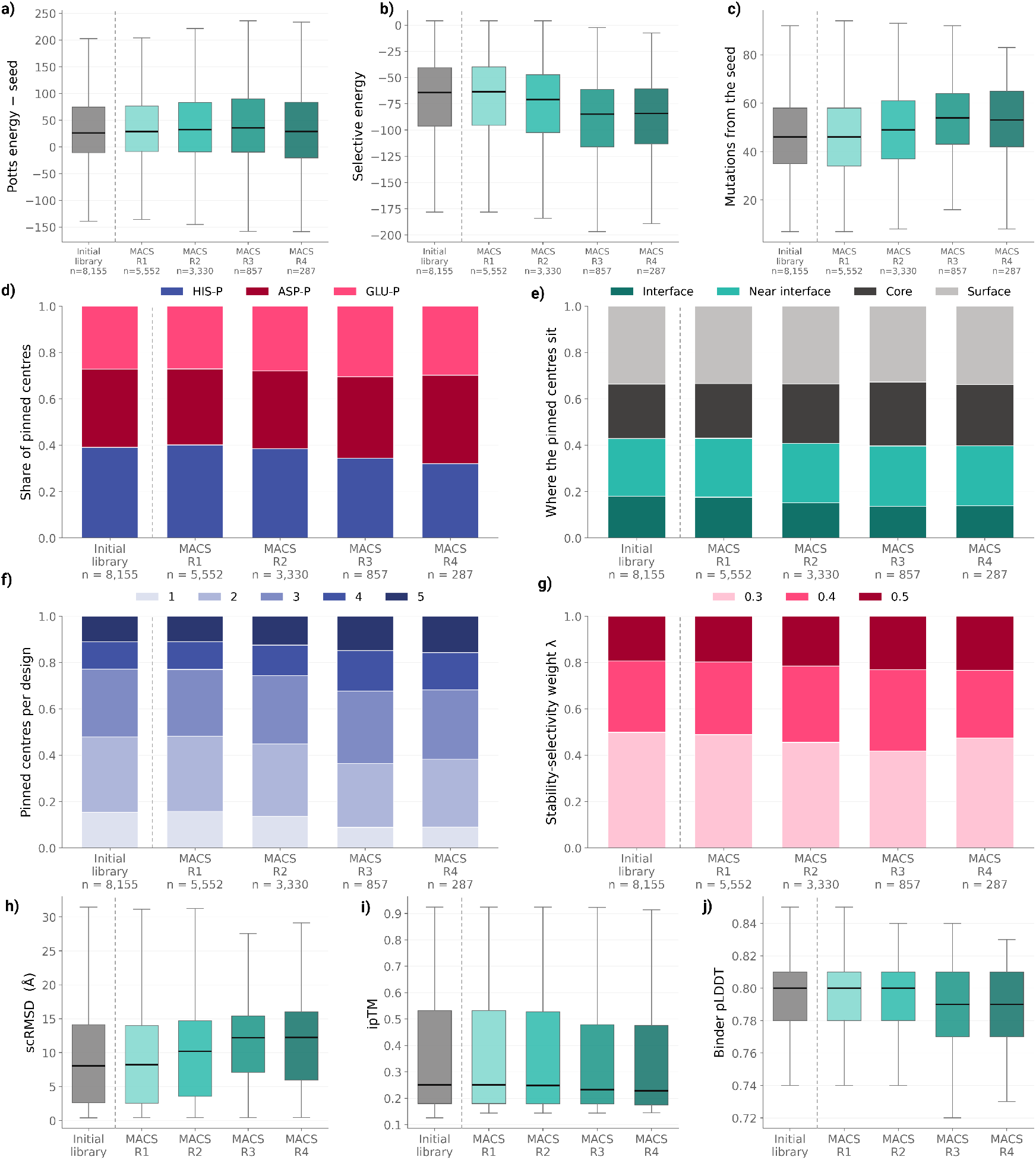
Properties of the PD-L1 library across the magnetic-sorting rounds. **a–c)** Distributions of the change in Potts energy relative to the seed sequence (**a**), selective energy summed over the pinned centers (**b**) and number of mutations from the seed sequence (**c**), for the initial library and the four rounds of magnetic-activated cell sorting. Boxes give the interquartile range and median and whiskers the 5th and 95th percentiles, here and in **h–j**. The dashed line separates the initial library from the selected pools, and *n* gives the number of designs detected in each pool. **d–g)** Composition of the same pools by the protonation state of the pinned centers (**d**), their structural location (**e**), the number of pinned centers per design (**f**) and the stability–selectivity weight *λ* used to generate them (**g**). **h–j)** Distributions of self-consistency RMSD (**h**), ipTM (**i**) and binder pLDDT (**j**) across the same pools. These structures were predicted after selection for analysis only; no structure-prediction filter was applied to the library at any stage.

**Fig. S17.**
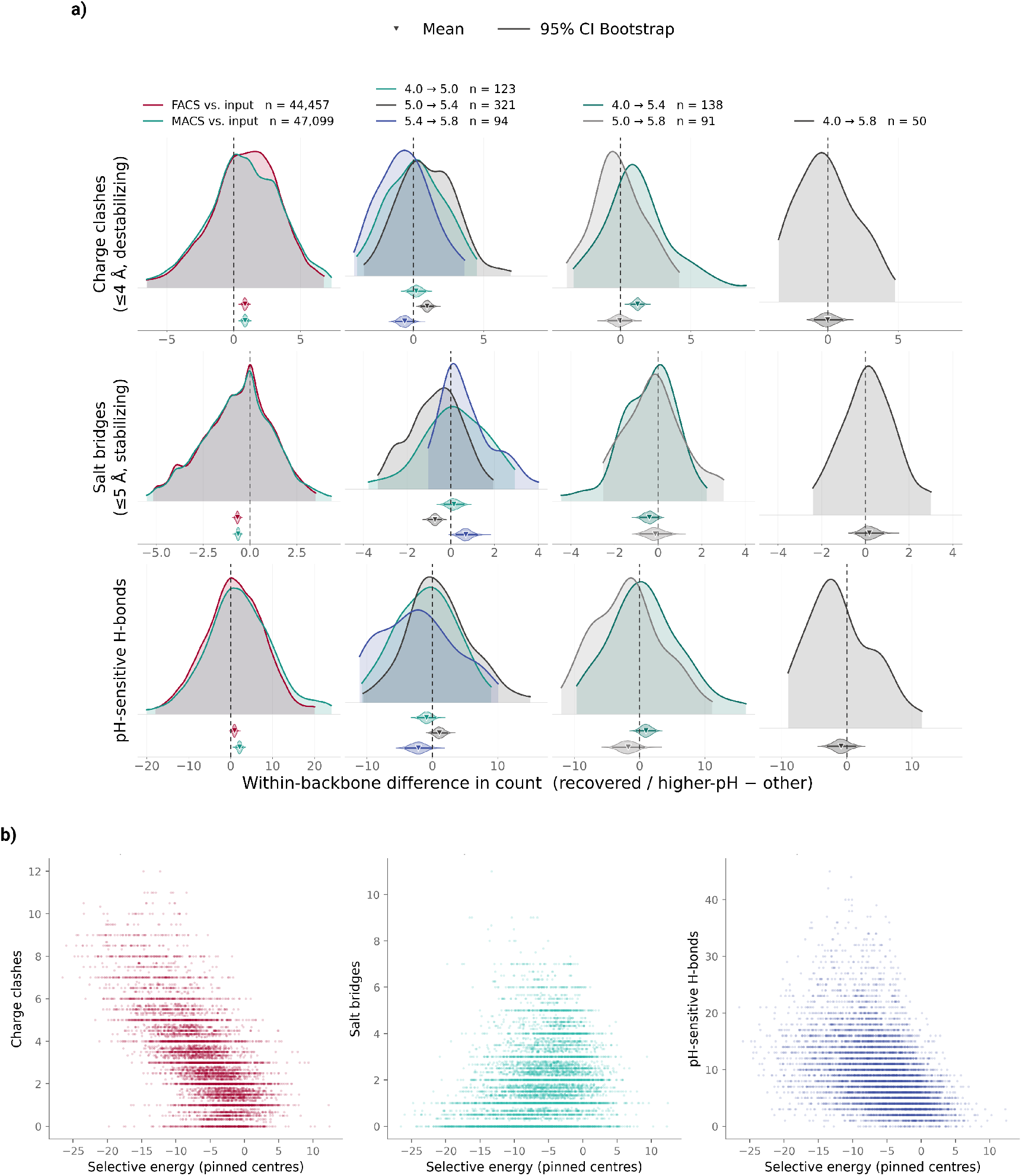
Charge interactions and pH-sensitive hydrogen bonds between matched backbone designs. **a)** Differences in the number of destabilising charge clashes (*≤*4 Å; top), stabilising salt bridges (*≤*5 Å; middle) and pH-sensitive hydrogen bonds (bottom) between designs sharing a backbone, computed as in Fig. 3f but for interaction counts rather than selective energy. The leftmost column compares recovered designs with the input library for the FACS and MACS tracks; the remaining columns compare same-backbone designs assigned to the indicated pH-response bins, with the difference taken as the design retained at the higher pH minus the design lost at the lower pH. Curves show paired differences, shaded distributions bootstrap estimates of the mean, and horizontal bars 95% confidence intervals; the dashed line marks zero and *n* is the number of matched pairs. **b) b)** Relationship between the selective energy at the pinned centers and the number of each interaction types. Each point is one design. Destabilising charge clashes (left, *≤*4 Å), stabilising salt bridges (middle) and pH-sensitive hydrogen bonds.

**Fig. S18.**
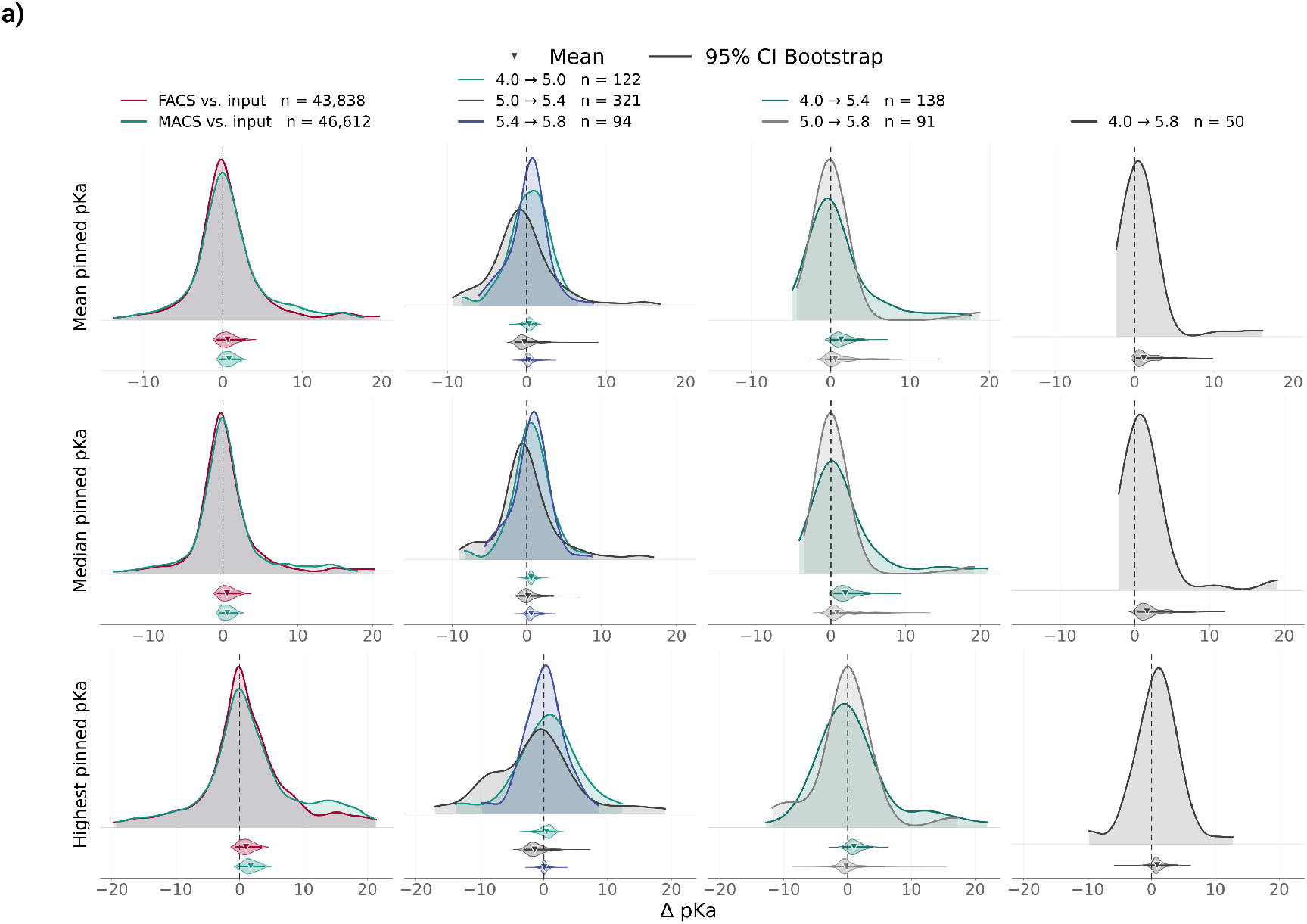
PROPKA3 p*K*_a_ at the pinned centers between matched backbone designs. Paired differences in PROPKA3-predicted p*K*_a_ at the pinned centers between designs sharing a backbone, computed as in Fig. 3f but for predicted p*K*_a_ rather than selective energy. Each design is summarised by the mean (top), median (middle) and highest (bottom) p*K*_a_ across its pinned centers. The leftmost column compares recovered designs with the input library for the FACS and MACS tracks. The remaining columns compare same-backbone designs in the indicated pH-response bins, with the difference taken as the design retained at the higher pH minus the design lost at the lower pH. Shaded distributions give bootstrap estimates of the mean and horizontal bars 95% confidence intervals, the dashed line marks zero and *n* is the number of matched pairs.

